# ANGPTL4 from adipose, but not liver, is responsible for regulating plasma triglyceride partitioning

**DOI:** 10.1101/2020.06.02.106815

**Authors:** Kathryn M. Spitler, Shwetha K. Shetty, Emily M. Cushing, Kelli L. Sylvers-Davie, Brandon S.J. Davies

## Abstract

Elevated plasma triglyceride levels are associated with metabolic disease. Angiopoietin-like protein 4 (ANGPTL4) regulates plasma triglyceride levels by inhibiting lipoprotein lipase (LPL). Our aim was to investigate the role of tissue-specific ANGPTL4 expression in the setting of high fat diet. Adipocyte- and hepatocyte-specific ANGPTL4 deficient mice were fed a high fat diet (60% kCal from fat) for either 12 weeks or 6 months. We performed plasma metabolic measurements, triglyceride clearance and uptake assays, LPL activity assays, and assessed glucose homeostasis. Mice lacking adipocyte ANGPTL4 recapitulated the triglyceride phenotypes of whole-body ANGPTL4 deficiency, whereas mice lacking hepatocyte ANGPTL4 had few triglyceride phenotypes. When fed a high fat diet (HFD), mice deficient in adipocyte ANGPTL4 gained more weight, had enhanced adipose LPL activity, and initially had improved glucose and insulin sensitivity. However, this improvement was largely lost after 6 months on HFD. Conversely, mice deficient in hepatocyte ANGPTL4 initially displayed no differences in glucose homeostasis, but began to manifest improved glucose tolerance after 6 months on HFD. We conclude that it is primarily adipocyte-derived ANGPTL4 that is responsible for regulating plasma triglyceride levels. Deficiency in adipocyte- or hepatocyte-derived ANGPTL4 may confer some protections against high fat diet induced dysregulation of glucose homeostasis.

## INTRODUCTION

Elevated plasma triglyceride levels have been implicated in the pathology of a variety of cardiovascular and metabolic diseases. Circulating triglycerides are hydrolyzed by lipoprotein lipase (LPL), releasing fatty acids that are taken up and utilized by tissues. Angiopoietin-like protein 4 (ANGPTL4), a secreted factor induced by fasting and expressed highly in adipose and liver, inhibits LPL activity and thereby regulates plasma triglyceride levels (1–5). Genetic loss of ANGPTL4 leads to decreased fasting plasma triglyceride levels in both humans and mice, and humans deficient in ANGPTL4 appear to be protected from cardiovascular disease (6–8). Moreover, a recent study found that human carriers of the common inactivating mutant E40K were protected against obesity-associated dyslipidemia (9). However, whether these protections are conferred by loss of ANGPTL4 in a specific tissue or require systemic loss of ANGPTL4 remains unclear.

In the last twenty years the work of many labs has led to development of a model where, during fasting, adipose-derived ANGPTL4 is rapidly induced and acts locally to inhibit adipose LPL activity and triglyceride uptake, diverting triglycerides to other tissues (3, 10–13). Recent studies using *Angptl4*^−/−^ or adipose-specific *Angptl4* knockout mice strongly support this model by showing that fasted mice lacking ANGPTL4 have increased uptake of triglycerides specifically in adipose tissue (14, 15). ANGPTL4 is also expressed in other metabolically active tissues, especially the liver. The role of ANGPTL4 in the liver remains unclear. Furthermore, the role of ANGPTL4 in mediating or alleviating metabolic disturbances induced by high-fat feeding has not been fully studied. Such studies have been hindered by the fact that *Angptl4*^−/−^ mice fed a high fat diet (HFD) develop a lethal intestinal injury and lymphatic nodule inflammation (16).

In this study, we sought to define the tissue-specific actions of ANGPTL4 more clearly. To this end, we generated adipocyte and hepatocyte-specific *Angptl4* knockout mice and studied their triglyceride and metabolic phenotypes when fed normal chow and high-fat diets. The effects of adipocyte and hepatocyte-specific ANGPTL4-deficiency on triglyceride and glucose homeostasis were determined after 12 weeks of high-fat feeding, as well as after 6 months of high-fat feeding to model a chronic state of obesity.

## RESULTS

### Effects of tissue-specific ANGPTL4 deficiency on plasma TG levels and TG uptake into tissues

To examine the tissue-specific roles of ANGPTL4, we generated adipocyte- and hepatocyte-specific *Angptl4* knockout mice. First, we generated *Angptl4*-floxed mice (*Angptl4*^fl/fl^) by utilizing CRISPR/Cas9 to insert LoxP sites in the introns between exons 1 and 2 and between exons 3 and 4 (**Figure 1A**). Recombination between LoxP sites generates a frame-shift resulting in a truncated (145 aa) protein in which the final 35 amino acids do not match the native sequence. Adipocyte-specific (*Angptl4*^AdipoKO^) and hepatocyte-specific (*Angptl4*^LivKO^) mice were generated by crossing *Angptl4*^fl/fl^ mice with adiponectin-Cre and albumin-Cre mice, respectively. To assess the impact of Cre-mediated recombination on *Angptl4* expression, we performed qPCR with 7 different primer sets on brown adipose tissue (BAT) from *Angptl4*^AdipoKO^ mice and on liver tissue from *Angptl4*^LivKO^ mice (**Supplemental Figure 1A**). Compared to Cre negative controls, significantly reduced expression was observed with all 7 primer sets (**Supplemental Figure 1B,C**). qPCR signal was further reduced when using primer sets targeting Exons 2 and 3, the exons removed by recombination (**Supplemental Figure 1B,C**). Together these data implied that Cre-mediated recombination was successful and that the recombined gene resulted in greatly reduced RNA expression of *Angptl4*.

**Figure 1.**
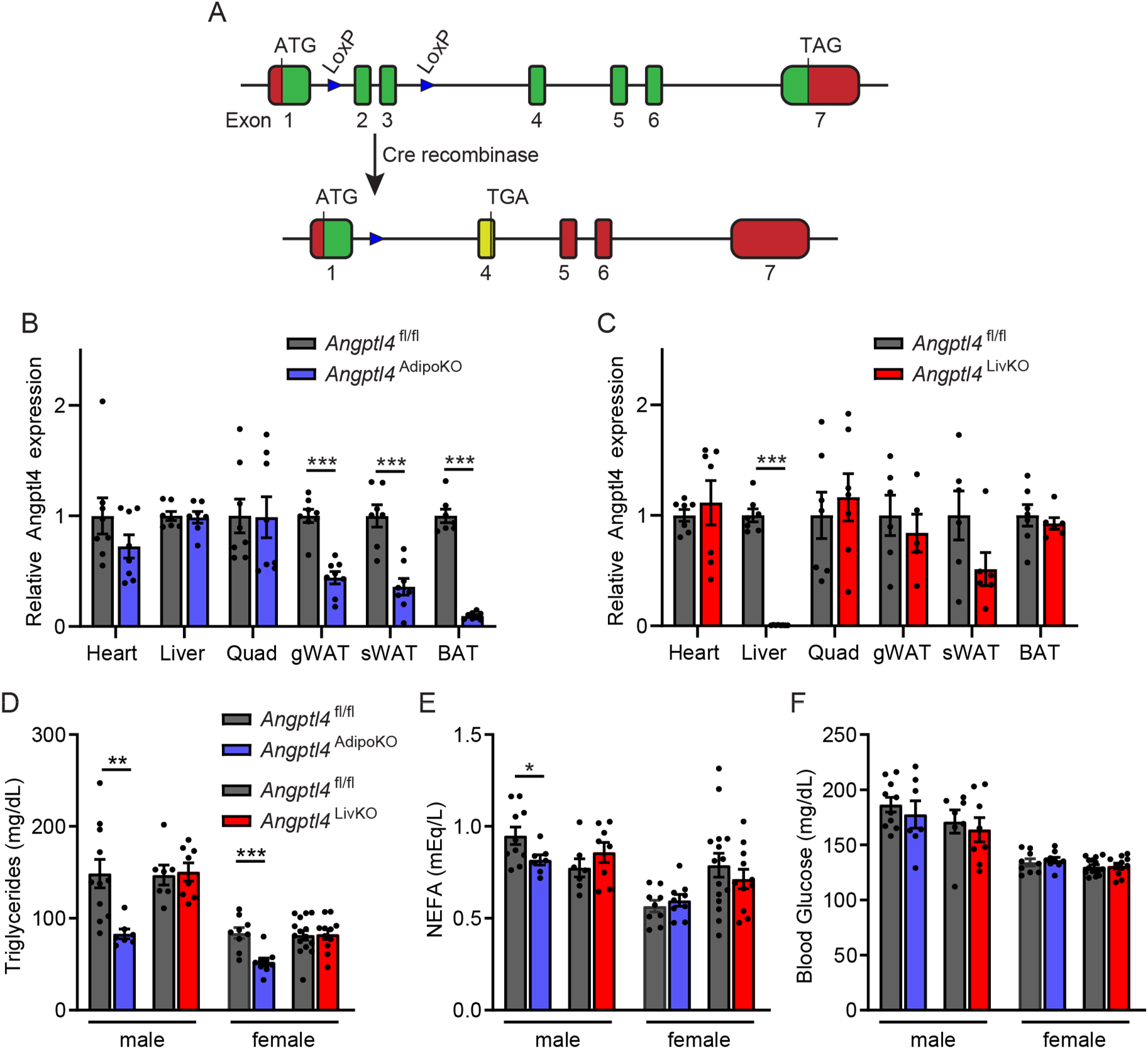
Generation and characterization of mice with adipose or liver-specific deletion of *Angptl4*. **A)** Schematic illustration of the lox-P sites in the *Angptl4*^fl/fl^ allele and genomic rearrangement that results from Cre-mediated recombination. **B and C)** mRNA expression of *Angptl4* in liver, heart, quadriceps muscle (quad), gonadal white adipose tissue (gWAT), subcutaneous white adipose tissue (sWAT), and brown adipose tissues (BAT) from 8-12 week old female *Angptl4*^fl/fl^, *Angptl4*^AdipoKO^ (B) and *Angptl4*^LivKO^ (C) mice following a 6 h fast (mean ± SEM; n=5-8). **D–F)** Fasting plasma triglyceride (D), non-esterified fatty acid (E) and blood glucose (F) levels in 8-12 week male and female *Angptl4*^fl/fl^, *Angptl4*^AdipoKO^, and *Angptl4*^LivKO^ mice following a 6 h fast (mean ± SEM; n=7-15). Bars represent mean ± SEM (n=7-11). **p<0.01, ***p<0.001. vs. *Angptl4*^fl/fl^ mice by student t-test (B-E).

Ideally, levels of ANGPTL4 protein would be assessed to ensure knockout of ANGPTL4, however no currently available antibodies are specific for mouse ANGPTL4. To assess if our recombined allele would still make protein, we generated a plasmid construct mimicking the predicted protein product of our *Angptl4* flox allele after Cre-mediated recombination (flox). We also generated a plasmid construct mimicking the post-recombination protein product of the *Angptl4* flox allele generated by the KOMP mouse repository and used in other studies of the tissue-specific actions of ANGPTL4 (14, 17, 18). 293T cells were transfected with these constructs or a construct expressing full-length wildtype mouse ANGPTL4 (wt) and both cell lysate and the media were subjected to Western blot analysis. As expected, in cells transfected with the full-length construct (wt), we observed ANGPTL4 protein in both the cell lysate and secreted in the media (**Supplemental Figure 1D**). We observed no protein expression in either the lysate or the media from the construct representing our flox allele, suggesting that after Cre recombination our flox allele does not produce protein. Interestingly, we did observe substantial protein expression in the lysate from the construct representing the KOMP allele, though very little of this protein appeared to be secreted (**Supplemental Figure 1D**). When the conditioned media from these constructs were tested for their ability to inhibit LPL, neither the media from our allele, nor the media containing the small amount of secreted protein from the KOMP flox allele were able to inhibit LPL activity (**Supplemental Figure 1E**).

We examined *Angptl4* expression in metabolically active tissues in both adipocyte-specific and hepatocyte-specific *Angptl4*-deficient mice. As expected, expression of *Angptl4* in gonadal, subcutaneous, and brown adipose tissues (gWAT, sWAT, BAT) was significantly reduced in *Angptl4*^AdipoKO^ mice, while being preserved in other tissues (**Figure 1B**). Likewise, in *Angptl4*^LivKO^ mice, *Angptl4* expression was significantly reduced only in the liver (**Figure 1C**). Fasting plasma triglyceride levels were significantly lower in 8-week-old male and female *Angptl4*^AdipoKO^ mice compared to *Angptl4*^fl/fl^ mice (**Figure 1D**), whereas TG levels were not different in either male or female *Angptl4*^LivKO^ mice compared to *Angptl4*^fl/fl^ mice (**Figure 1D**). These results suggest that the lower plasma TG levels observed in *Angptl4* whole-body knockout mice (2, 15, 19) is due to the loss of adipocyte-derived ANGPTL4. We observed a decrease in fasting plasma free fatty acid levels in male, but not female *Angptl4*^AdipoKO^ mice compared to *Angptl4*^fl/fl^ mice, but no difference in free fatty acids levels in either male or female *Angptl4*^LivKO^ mice compared to *Angptl4*^fl/fl^ mice (**Figure 1E**). There were no genotype-specific differences in fasting blood glucose levels between groups (**Figure 1F**).

We previously observed increased plasma triglyceride clearance, increased adipose uptake of triglycerides, and increased adipose LPL activity in whole-body *Angptl4*^−/−^ mice (15). In that study we hypothesized that the increased clearance was due to ANGPTL4 deficiency specifically in the adipose tissue (15). To determine if this was indeed the case, we performed triglyceride clearance assays on female *Angptl4*^AdipoKO^ and *Angptl4*^LivKO^ mice using radiolabeled chylomicrons. We also assessed uptake of radiolabeled chylomicrons into specific tissues. After an intravenous injection of radiolabeled chylomicrons, *Angptl4*^AdipoKO^ mice cleared the radiolabeled TGs from the plasma faster than *Angptl4*^fl/fl^ mice (**Figure 2A**). The rate of radiolabeled clearance was also modestly increased in *Angptl4*^LivKO^ mice compared to *Angptl4*^fl/fl^ mice. (**Figure 2B**). Similar to what we had observed with whole-body *Angptl4*^−/−^ mice (15), *Angptl4*^AdipoKO^ female mice had increased radiolabel uptake into adipose depots compared to *Angptl4*^fl/fl^ mice (**Figure 2C**). No significant differences in tissue uptake were observed in female *Angptl4*^LivKO^ mice, other than a decrease in TG uptake in the quadriceps (**Figure 2D**). As with whole-body *Angptl4*^−/−^ mice, LPL activity was increased in white adipose tissue of female *Angptl4*^AdipoKO^ mice compared to *Angptl4*^fl/fl^ mice (**Figure 2E,G**). Additionally, there was a decrease in LPL activity in heart tissue of female *Angptl4*^AdipoKO^ mice compared to *Angptl4*^fl/fl^ mice. No differences in LPL activity were observed in any tissue from female *Angptl4*^LivKO^ mice compared to female *Angptl4*^fl/fl^ mice (**Figure 2F,H**). These data support our previous hypothesis that deficiency in adipose ANGPTL4, not liver ANGPTL4, is responsible for the increased triglyceride uptake into adipose tissue.

**Figure 2:**
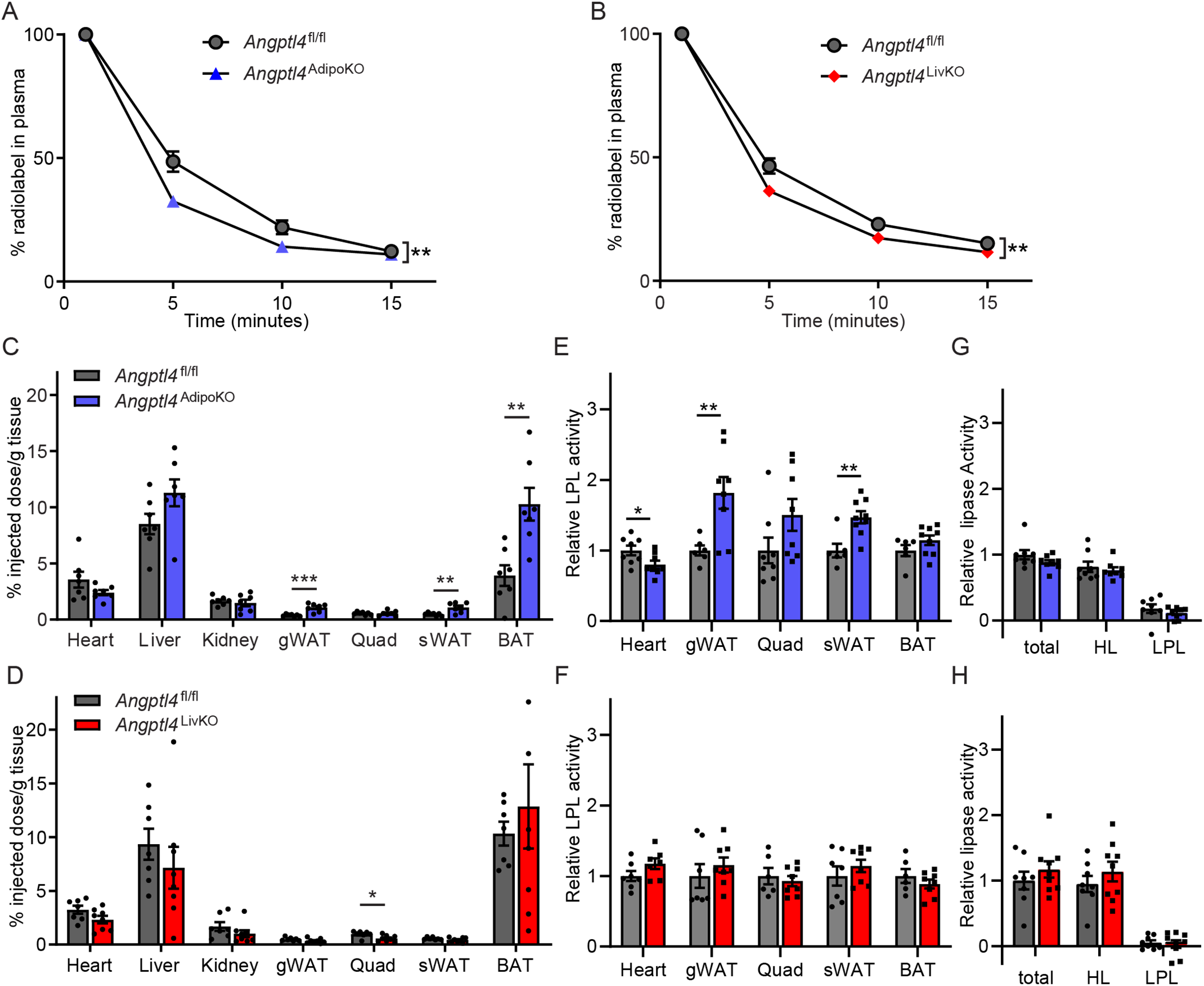
Chylomicron clearance and uptake and lipase activity in female *Angptl4*^AdipoKO^ and *Angptl4*^LivKO^ mice. **A–D)** Fasted (6 h) female *Angptl4*^AdipoKO^ (A,C) and *Angptl4*^LivKO^ (B,D) mice were injected intravenously with ^3^H-triglyceride-containing chylomicrons. **A and B)** Clearance of radiolabel from the plasma 1, 5, 10, and 15 min after injection. Points represent percentage of radiolabel remaining in the plasma at the indicated time points compared to the 1 min time point (mean±SEM; n=7-8). **C and D)** Uptake of radiolabel after 15 min (% injected dose/g tissue) into the indicated tissues (mean±SEM; n=7-8). **E-F)** Heart, gonadal white adipose tissue (gWAT), quadricep muscle (Quad), subcutaneous adipose tissue (sWAT), and brown adipose (BAT) tissue from fasted (6 h) female *Angptl4*^AdipoKO^ (E) and *Angptl4*^LivKO^ (F) mice were harvested and lipase activity was measured (n=6-9/group). **G and H**) Liver was harvested from fasted (6 h) female *Angptl4*^AdipoKO^ (G) and *Angptl4*^LivKO^ (H) mice (n=8-9/group). Lipase activity was measured in the presence or absence of 1M-NaCl to distinguish between hepatic and lipoprotein lipase. Bars show relative lipase activity in each tissue normalized to *Angptl4*^fl/fl^ (mean±SEM). *p<0.05, **p<0.01, ***p<0.001 vs *Angptl4*^fl/fl^ mice for individual genotype-specific differences by repeated measures ANOVA (A,B) or t-test analysis (C-H).

### Effects of high-fat feeding on body phenotypes and plasma parameters in tissue-specific ANGPTL4-deficient mice

Given the importance of ANGPTL4 in directing triglycerides away from adipose tissue (15), we sought to determine how high-fat feeding would alter metabolic phenotypes in *Angptl4*^AdipoKO^, *Angptl4*^LivKO^, and *Angptl4*^fl/fl^ mice. We randomly assigned *Angptl4*^AdipoKO^, *Angptl4*^LivKO^ and *Angptl4*^fl/fl^ mice at 8 weeks of age to a normal chow (NCD) or high fat diet (HFD; 60% kCal/fat) and fed them the respective diet for 12 weeks. On HFD, body weight gain was similar between all groups (**Figure 3A,B**); however, in this cohort, *Angptl4*^LivKO^ mice fed a normal chow diet gained slightly less weight than *Angptl4*^fl/fl^ mice (**Figure 3B**). Adiposity, as determined by NMR, was greater in all mice fed a HFD compared to mice fed a NCD (**Figure 3C,D**). Despite no differences in body weight, *Angptl4*^*AdipoKO*^ mice had significantly more total fat mass compared to *Angptl4*^fl/fl^ mice when fed a NCD (**Figure 3C**). This genotype-specific difference in fat mass was not seen in HFD-fed *Angptl4*^*AdipoKO*^ mice, nor were any genotype-specific differences in fat mass observed in *Angptl4*^LivKO^ mice (**Figure 3C,D**). Moreover, no genotype-specific differences in specific tissue weights were observed in either *Angptl4*^AdipoKO^ or *Angptl4*^LivKO^ mice (**Supplemental Figure 2**).

**Figure 3.**
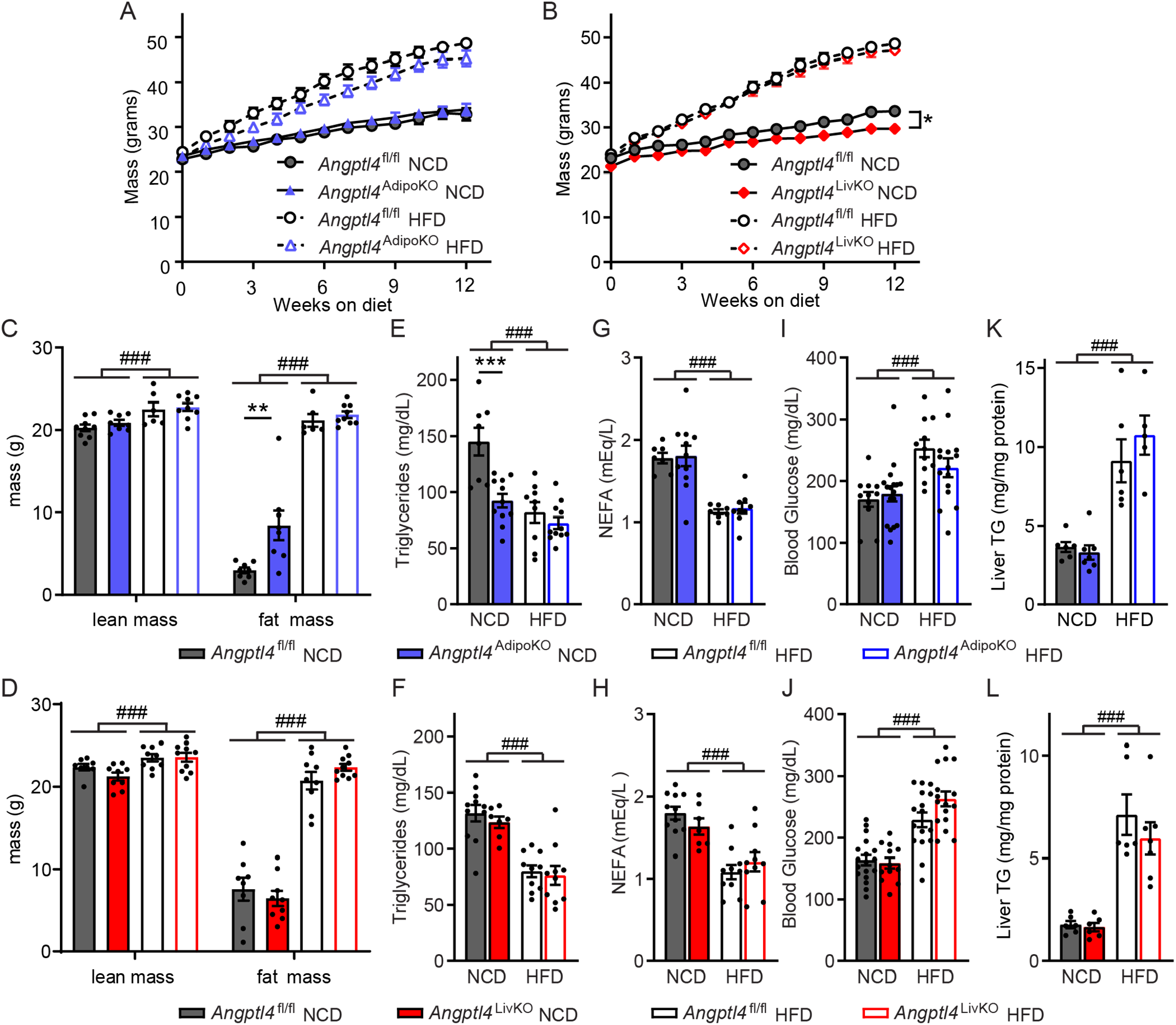
Body weights, fat mass, and metabolic phenotypes of *Angptl4*^AdipoKO^ and *Angptl4*^LivKO^ mice. **A and B)** Weekly body weights of male *Angptl4*^AdipoKO^ (A) and *Angptl4*^LivKO^ (B) mice fed either a normal chow diet (NCD) or a high fat diet (HFD; 60% by kCal) for 12 weeks starting at 8 weeks of age (mean±SEM; n=12-20/group). *p<0.05 by repeated measures ANOVA. **C and D)** Lean muscle and fat mass of male *Angptl4*^AdipoKO^ (C) and *Angptl4*^LivKO^ (D) mice as measured by NMR after 12 weeks on diet (mean±SEM; n=6-9/group). Plasma triglycerides **(E,F)**, plasma non-esterified fatty acids **(G,H)**, blood glucose **(I,J),** and liver triglycerides **(K,L)** of fasted (6 h) male *Angptl4*^AdipoKO^ and *Angptl4*^LivKO^ mice at the end of 12 weeks of either normal chow diet (NCD) or high fat diet (HFD; mean±SEM; n=8-22 for E-J and n=4-7/group for K,L). ###p<0.001 for dietary differences by two-way ANOVA. *p<0.05, **p<0.01, ***p<0.001 for individual genotype-specific differences by multiple comparison after two-way ANOVA (Tukey correction).

Tissue expression of *Angptl4* was influenced both by genotype and by diet. As expected, expression of *Angptl4* in gonadal, subcutaneous, and brown adipose tissues (eWAT, sWAT, BAT) was much lower in *Angptl4*^AdipoKO^ mice compared to *Angptl4*^fl/fl^ mice (**Supplemental Figure 3A**). Surprisingly, *Angptl4*^AdipoKO^ mice on NCD had greatly reduced *Angptl4* expression in the heart compared to *Angptl4*^fl/fl^ mice (**Supplemental Figure 3A**), suggesting that loss of ANGPTL4 in the adipose tissue had a systemic effect leading to downregulation of ANGPTL4 in other tissues. In *Angptl4*^LivKO^ mice, expression of *Angptl4* was reduced only in the liver compared to *Angptl4*^fl/fl^ mice (**Supplemental Figure 3B**). Interestingly, HFD-alone significantly decreased *Angptl4* expression in several tissues, including the heart, liver, and white adipose tissue (**Supplemental Figure 3A,B**). These data indicate that high fat feeding can regulate expression of *Angptl4*.

As before, *Angptl4*^AdipoKO^ mice, but not *Angptl4*^LivKO^ mice fed a NCD had significantly lower fasting plasma TG levels compared to *Angptl4*^fl/fl^ mice (**Figure 3E,F**). 12 weeks of HFD feeding lowered fasting TG levels in all groups and mostly negated the differences in plasma TG levels between *Angptl4*^AdipoKO^ and *Angptl4*^fl/fl^ mice (**Figure 3E,F**). HFD feeding for 12 weeks decreased plasma non-esterified fatty acid levels and increased plasma glucose and liver triglyceride levels in all groups, but there were no genotype specific differences on either NCD or HFD. (**Figure 3G-L**).

As changes in energy expenditure play a role in the pathogenesis of obesity, we asked if either hepatocyte- or adipocyte-specific loss of ANGPTL4 would lead to changes in energy expenditure, oxygen consumption (VO_2_), carbon dioxide production (VCO_2_), or respiratory quotient (RQ; VCO_2_/VO_2_). Using the Promethion metabolic caging system, we observed no differences in energy expenditure, VO_2_, VCO_2_, or RQ in *Angptl4*^AdipoKO^ and *Angptl4*^LivKO^ mice on NCD compared to *Angptl4*^fl/fl^ mice (**Supplemental Figure 4**). Interestingly, compared to floxed controls, increased energy expenditure was observed in the *Angptl4*^LivKO^ mice fed a HFD during both light and dark cycles. Both VO_2_ and VCO_2_ were increased in HFD-fed *Angptl4*^*LivKO*^ mice, while the RQ was unchanged. No differences were observed between *Angptl4*^AdipoKO^ and *Angptl4*^fl/fl^ mice on HFD (**Supplemental Figure 4**).

### Effects of high-fat feeding on TG uptake and LPL activity in tissue-specific *Angptl4* KO mice

To assess if high-fat diet feeding altered triglyceride partitioning in an ANGPTL4-dependent manner, we performed plasma triglyceride clearance and tissue uptake assays on male *Angptl4*^AdipoKO^ and *Angptl4*^LivKO^ mice at the end of 12 weeks of either NCD or HFD feeding. On a NCD, *Angptl4*^AdipoKO^ mice cleared the radiolabeled TGs from the plasma faster than *Angptl4*^fl/fl^ mice (**Figure 4A**), but no significant difference in triglyceride clearance was observed on a NCD for *Angptl4*^LivKO^ mice compared to *Angptl4*^fl/fl^ mice (**Figure 4B**). Mice fed a HFD had faster clearance of plasma triglycerides than those fed a NCD, but on HFD there were no genotype specific differences in clearance rates (**Figure 4A,B**). Consistent with our observations in female mice, on NCD *Angptl4*^AdipoKO^ male mice had increased radiolabel uptake into epididymal and subcutaneous adipose depots compared to floxed controls (**Figure 4C**). Interestingly, on HFD, the differences in radiolabel uptake into adipose tissues between *Angptl4*^AdipoKO^ mice and *Angptl4*^fl/fl^ littermates were reduced (**Figure 4C**). No significant differences in tissue uptake were observed in male *Angptl4*^LivKO^ mice fed either a NCD or HFD compared to *Angptl4*^fl/fl^ mice (**Figure 4D**). These data again support the idea that on a NCD, deficiency in adipose ANGPTL4 results in increased triglyceride uptake into adipose tissue, but also suggest that this difference decreases in the face of high-fat feeding.

**Figure 4.**
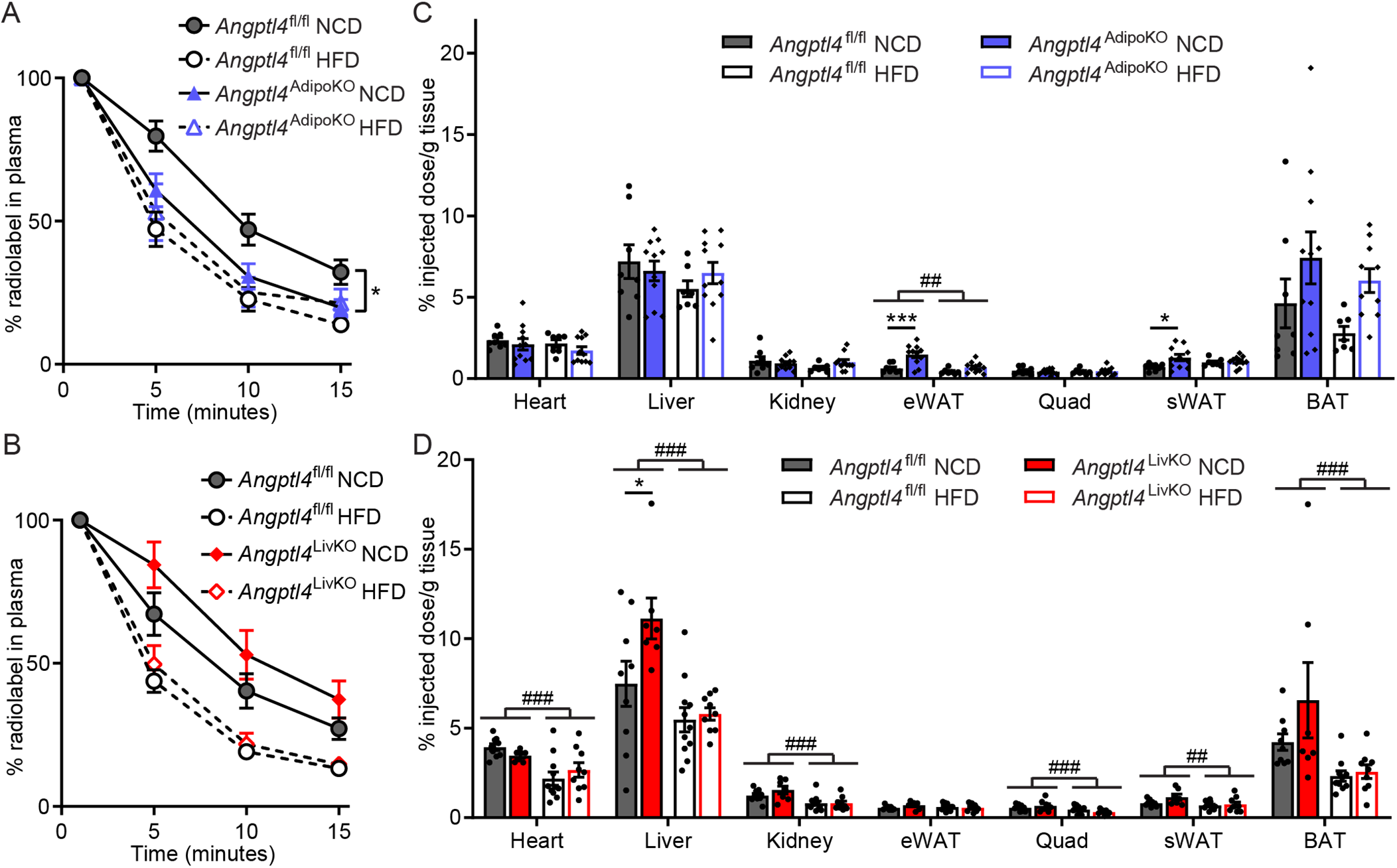
Chylomicron clearance and uptake and lipase activity in *Angptl4*^AdipoKO^ and *Angptl4*^LivKO^ mice. At the conclusion of 12 weeks of either normal chow diet (NCD) or high fat diet (HFD) *Angptl4*^AdipoKO^ (A,C) and *Angptl4*^LivKO^ (B,D) male mice (n=7-10/group) were fasted (6 h) and injected intravenously with ^3^H-triglyceride containing chylomicrons. **A and B)** Clearance of radiolabel from the plasma 1, 5, 10, and 15 minutes after injection. Points represent percentage of radiolabel remaining in the plasma at the indicated time points compared to the 1-minute time point (mean±SEM). *p<0.05 by repeated measures ANOVA. **C and D)** Uptake of radiolabel (% injected dose/g tissue) into the indicated tissues after 15 min (mean±SEM). ##p<0.01, ###p<0.001 for dietary differences by two-way ANOVA. *p<0.05, ***p<0.001 for individual genotype-specific differences by multiple comparison after two-way ANOVA (Tukey correction).

To elucidate if the increased uptake of triglyceride-derived fatty acids into adipose tissue in *Angptl4*^AdipoKO^ mice was the result of increased LPL activity we measured tissue lipase activity. LPL activity was greater in the gonadal adipose tissue of *Angptl4*^AdipoKO^ mice on either NCD or HFD compared to *Angptl4*^fl/fl^ mice (**Figure 5A**). There were no genotype specific differences in tissue LPL activity between *Angptl4*^LivKO^ mice and *Angptl4*^fl/fl^ mice (**Figure 5B**). HFD itself altered LPL activity in some tissues, increasing activity in quadriceps and slightly decreasing activity in epididymal white adipose tissue in both cohorts (**Figure 5A,B**). We also assessed if loss of ANGPTL4 resulted in changes in liver lipase activity. As expected, LPL activity in the liver was low compared to total lipase activity. We found no genotype specific differences in total lipase, hepatic lipase, or LPL activity on either diet in the *Angptl4*^AdipoKO^ mice compared to *Angptl4*^fl/fl^ mice (**Figure C,E**). While no genotype specific differences were found in *Angptl4*^LivKO^ mice in total lipase, hepatic lipase or LPL activity on NCD, there was a decrease in LPL activity in the *Angptl4*^LivKO^ mice on HFD compared to *Angptl4*^fl/fl^ mice (**Figure 5D,F**). We also measured tissue gene expression of *Lpl* and found no genotype-specific differences (**Supplemental Figure 5**). Together, TG clearance and LPL activity in *Angptl4*^*AdipoKO*^ mice closely resemble those we have previously reported in whole-body ANGPTL4 knockout mice (15), supporting the idea that the lack of adipose-derived ANGPTL4 is the primary driver of the increased adipose LPL activity and TG clearance in those mice.

**Figure 5.**
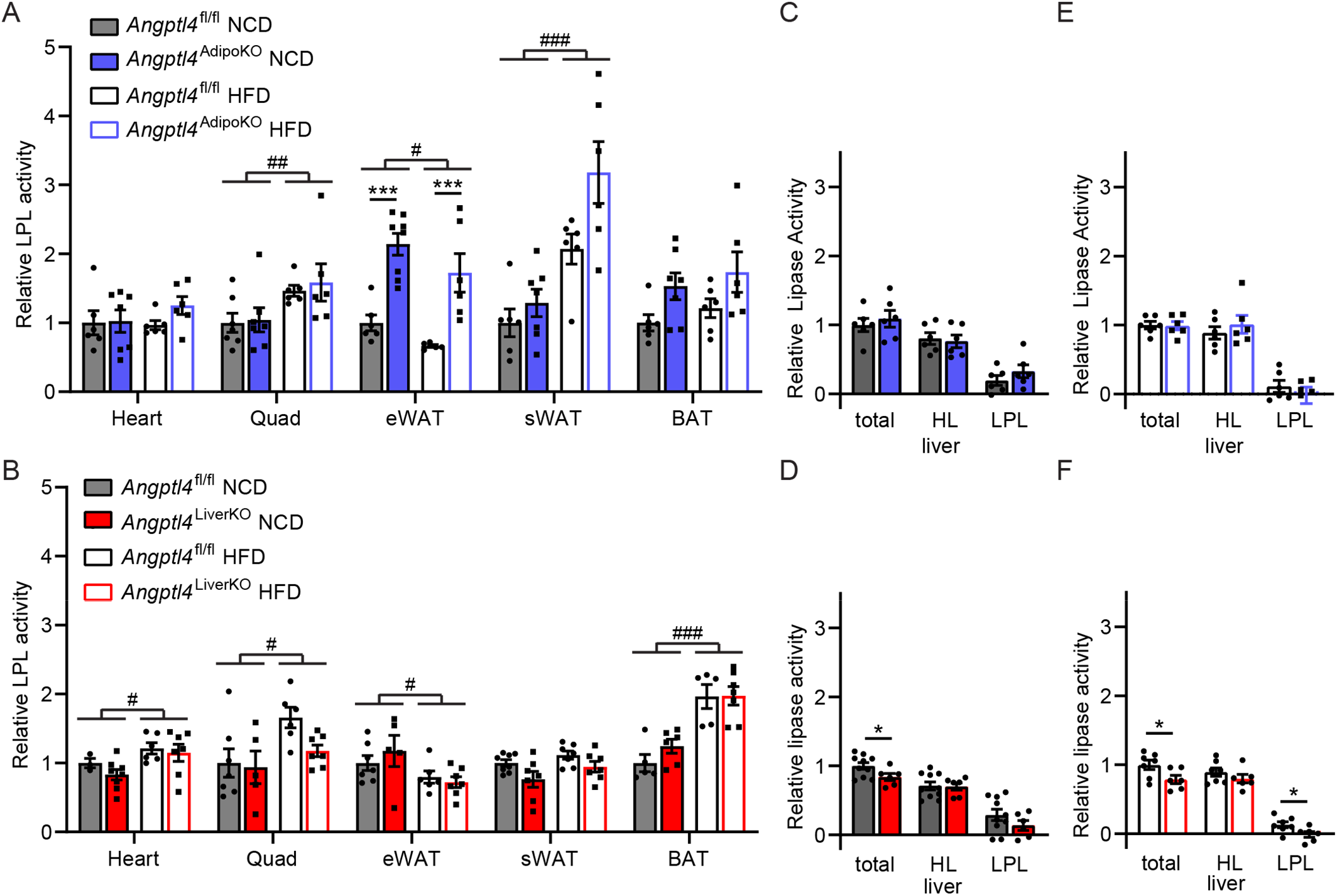
Lipase activity in *Angptl4*^AdipoKO^ and *Angptl4*^LivKO^ mice. **A and B)** Heart, epididymal adipose tissue (eWAT), quadricep muscle (Quad), subcutaneous adipose tissue (sWAT), and brown adipose (BAT) tissue from fasted (6 h) male *Angptl4*^AdipoKO^ (A) and *Angptl4*^LivKO^ (B) mice were harvested and lipase activity was measured (n=3-7/group). **C–F)** Liver was harvested from fasted (6 h) male *Angptl4*^AdipoKO^ (C, NCD groups and E, HFD groups) and *Angptl4*^LivKO^ (D, NCD groups and F, HFD groups) mice (n=4-7/group). Lipase activity was measured in the presence or absence of 1M NaCl to distinguish between hepatic and lipoprotein lipase. Bars show relative lipase activity in each tissue normalized to *Angptl4*^fl/fl^ (mean±SEM). #p<0.05, ##p<0.01, ###p<0.001 for dietary differences by two-way ANOVA. *p<0.05, ***p<0.001 for individual genotype-specific differences by multiple comparison after two-way ANOVA (Tukey correction).

### Effects of high-fat feeding on glucose tolerance and insulin sensitivity in tissue-specific ANGPTL4-deficient mice

A previous report found that an independent strain of adipocyte-specific ANGPTL4 knockout mice had improved glucose tolerance compared to wild-type mice after being fed a HFD for 4 weeks (14). To assess if the loss of adipocyte or hepatocyte derived ANGPTL4 altered glucose metabolism in our mice, we performed glucose (GTT) and insulin tolerance (ITT) tests. On a normal chow diet, no differences were observed in glucose tolerance or insulin sensitivity in either *Angptl4*^AdipoKO^ or *Angptl4*^LivKO^ mice compared to floxed controls (**Figure 6**). However, glucose tolerance and insulin sensitivity were markedly improved in HFD-fed *Angptl4*^AdipoKO^ mice compared to HFD-fed floxed controls (**Figure 6A,C**). The loss of hepatocyte-derived ANGPTL4 resulted in no improvements in glucose tolerance or insulin sensitivity on either diet (**Figure 6B,D**). These data support a protective role for loss of adipose-derived ANGPTL4 on systemic glucose homeostasis. Chronic inflammation has been linked with insulin resistance during obesity and type 2 diabetes (20). To test whether the changes we observed in glucose tolerance and insulin sensitivity were due to changes in tissue inflammation we measured tissue expression (liver, eWAT, sWAT, and BAT) of inflammatory markers C-C Motif Chemokine Ligand 2 (*Ccl2), Cd68*, and tumor necrosis factor α (*Tnfα*). As expected, there was an increase in inflammation in all tested tissues of HFD-fed mice. However, no major genotype specific differences were observed (**Supplemental Figure 6**).

**Figure 6.**
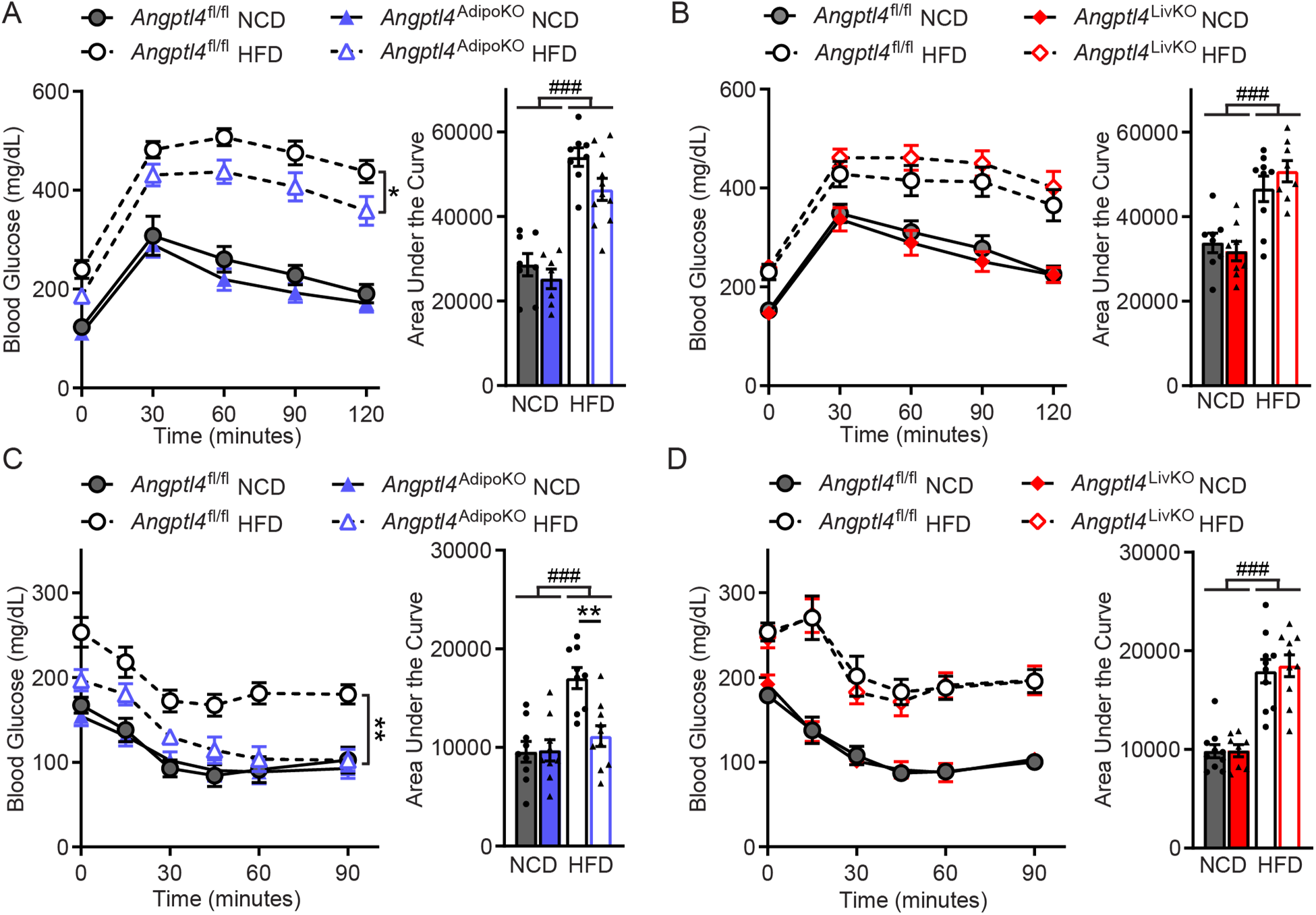
Glucose tolerance and insulin sensitivity of *Angptl4*^AdipoKO^ and *Angptl4*^LivKO^ mice. **A and B)** Glucose tolerance tests were performed on fasted (6 h) male *Angptl4*^AdipoKO^ (A) and *Angptl4*^LivKO^ (B) mice after 11 weeks of either a normal chow diet (NCD) or high fat diet (HFD). Mice were injected with glucose (2 g/kg for NCD; 1.3 g/kg HFD) and blood glucose concentrations were measured over 2 h. Points represent glucose levels (mean±SEM; n=7-11) at each respective time point. Bar graphs represent area under the curve (mean±SEM) for all time points. *p<0.05 by repeated measures ANOVA. **C and D)** Insulin tolerance tests were performed on fasted (4 h) male *Angptl4*^AdipoKO^ (C) and *Angptl4*^LivKO^ (D) mice after 12 weeks of either a normal chow diet (NCD) or high fat diet (HFD). Mice were injected with 0.75 U/ml of human insulin (Humalin-R) and blood glucose concentrations were measured over 90 min. Points represent glucose levels (mean±SEM; n=7-11) at each respective time point. Bar graphs represent area under the curve (mean±SEM) for all time points. ###p<0.001 for dietary differences by two-way ANOVA. **p<0.01 for individual genotype-specific differences by repeated measures ANOVA.

### Effects of tissue specific ANGPTL4 deficiency in mice after a chronic high-fat diet

To better characterize the role of ANGPTL4 in the setting of chronic obesity, we fed *Angptl4*^AdipoKO^, *Angptl4*^LivKO^, and *Angptl4*^fl/fl^ mice a NCD or HFD (60% kCal/fat) for 6 months. Initially, there were no genotype-specific differences in weight gain between any of the diet groups (**Figure 7A,B**). However, after 18 weeks on diet, the body weight curves began to diverge, with *Angptl4*^*AdipoKO*^ mice gaining more weight than littermate *Angptl4*^*fl/fl*^ mice on both a NCD or and a HFD (**Figure 7A**). At the end of 6 months of high-fat feeding, *Angptl4*^LivKO^ mice weighed slightly more than their high-fat fed *Angptl4*^*fl/fl*^ controls (**Figure 7B**). Adiposity, as determined by NMR, was greater in all mice fed a HFD compared to mice fed a NCD, and there was a significant increase in total fat mass in the high fat fed *Angptl4*^*AdipoKO*^ mice compared to high fat fed *Angptl4*^*fl/fl*^ controls (**Figure 7C,D**). There were no genotype specific differences in tissue weights (**Supplemental Figure 7**). As expected, expression of *Angptl4* in the gonadal, subcutaneous and brown adipose tissues (eWAT, sWAT, BAT) of *Angptl4*^AdipoKO^ mice and in the liver of *Angptl4*^LivKO^ mice were much lower than in *Angptl4*^fl/fl^ littermates (**Supplemental Figure 8**). Similar to what we observed after 12 weeks on diet, expression of *Angptl4* was reduced in *Angptl4*^*AdipoKO*^ mice in the heart and quadricep muscle compared to *Angptl4^fl/fl^*. Also similar to what we observed after 12 weeks of high-fat feeding, *Angptl4* expression was reduced in heart, liver, and white adipose tissue of Angptl4^fl/fl^ mice fed a HFD for 6 months, again supporting the idea that *Angptl4* expression in these tissues is regulated by diet.

**Figure 7.**
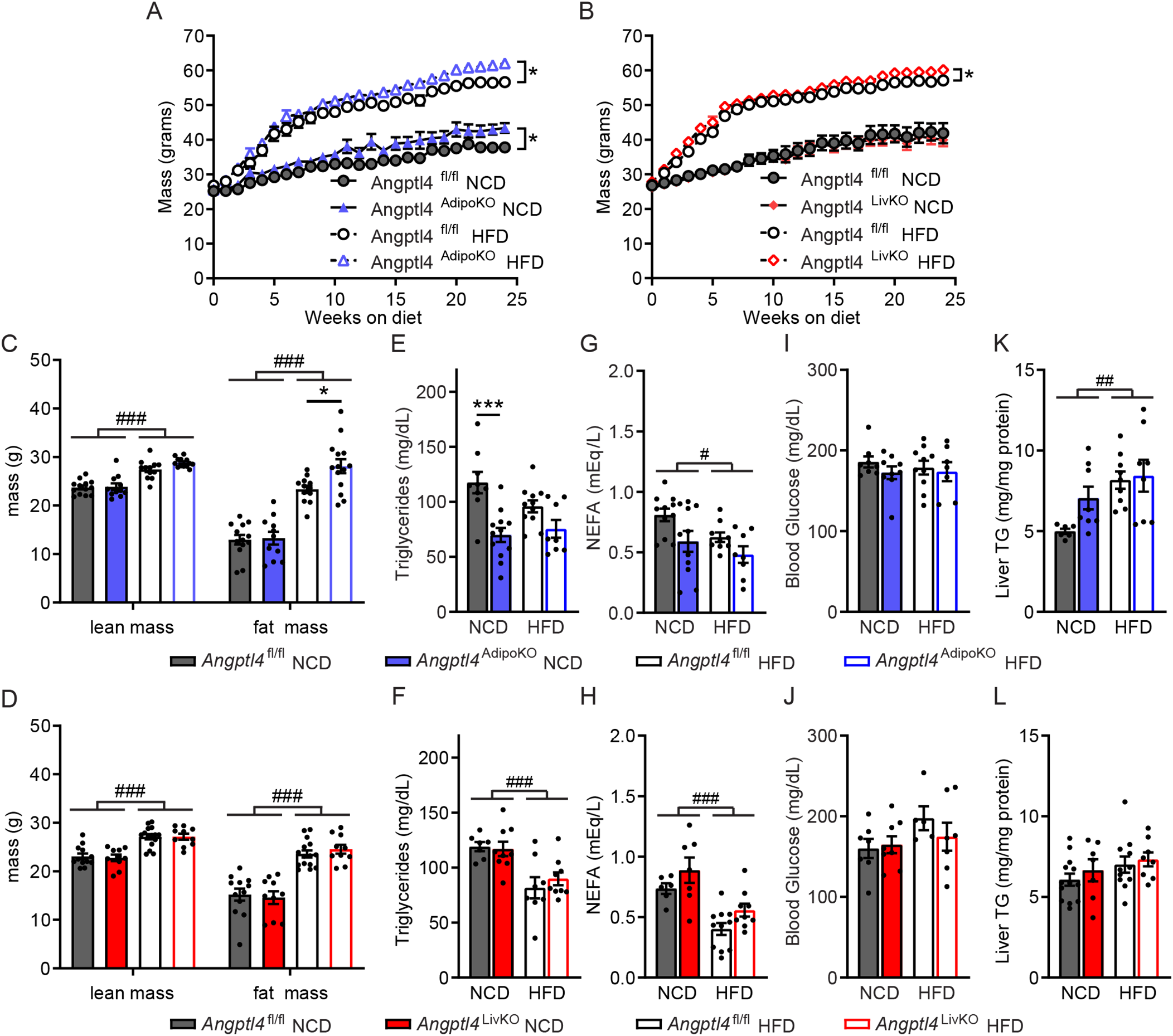
Body weights, fat mass, and metabolic phenotypes of *Angptl4*^AdipoKO^ and *Angptl4*^LivKO^ mice after chronic high-fat feeding. **A and B)** Weekly body weights of male *Angptl4*^AdipoKO^ (A) and *Angptl4*^LivKO^ (B) mice fed either a normal chow diet (NCD) or high fat diet (HFD) for 6 months starting at 8 weeks of age (mean±SEM; n=15-20/group). *p<0.05 by repeated measures ANOVA. **C and D)** Lean muscle mass and fat mass of male *Angptl4*^AdipoKO^ (C) and *Angptl4*^LivKO^ (D) mice as measured by NMR after 24 weeks on diet. (mean±SEM; n=10-13/group). Plasma triglycerides **(E,F)**, plasma non-esterified fatty acids **(G,H)**, blood glucose **(I,J),** and liver triglycerides **(K,L)** of fasted (6 h) male *Angptl4*^AdipoKO^ and *Angptl4*^LivKO^ mice after 6 months of either normal chow diet (NCD) or a high fat diet (HFD) feeding (mean±SEM; n=7-12/group). #p<0.05, ##p<0.01, ###p<0.001 for dietary differences by two-way ANOVA. *p<0.05,***p<0.001 for individual genotype-specific differences by multiple comparison after two-way ANOVA (Tukey correction).

After 6 months on diet, *Angptl4*^AdipoKO^ mice fed a NCD continued to have significantly lower fasting plasma triglyceride levels compared to *Angptl4*^*fl/fl*^ mice, but on a HFD these differences were not significant (**Figure 7E**). As before, no genotype-specific differences in plasma TG levels were observed between *Angptl4*^*LivKO*^ mice and *Angptl4*^*fl/fl*^ controls (**Figure 7F**). At this age, ANGPTL4-deficiency in either adipocytes or hepatocytes had no significant effect on plasma non-esterified fatty acids, blood glucose, or liver triglycerides levels (**Figure 7G-L**). Interestingly, although HFD-fed mice had lower fasting plasma non-esterified free fatty acid levels than NCD-fed mice, at this time point, the diet induced differences in blood glucose and liver TG mostly disappeared (**Figure 7G-L**). To assess liver health in our mice we measured plasma activity of alanine aminotransferase (ALT) and aspartate aminotransferase (AST). All mice fed a HFD for 6 months had elevated AST and ALT activity compared to NCD fed mice, but no genotype specific differences were observed (**Supplemental Figure 9A-D**). We also measured inflammatory protein amyloid A in the plasma of our mice. Although HFD-fed mice had an increase in plasma amyloid A there were no genotype-specific differences (**Supplemental Figure 9E,F**). Livers of mice fed a HFD appeared larger than those from mice fed NCD, but again no genotype specific differences were observed (**Supplemental Figure 9G,H**).

After 6 months on diet, *Angptl4*^AdipoKO^ mice continued to clear radiolabeled triglycerides from the plasma faster than littermate *Angptl4*^fl/fl^ mice on NCD (**Figure 8A**). *Angptl4*^AdipoKO^ mice also cleared triglycerides faster when fed a HFD, though the difference was less pronounced (**Figure 8A**). No genotype-specific difference in radiolabel clearance was observed in *Angptl4*^LivKO^ mice compared to littermate *Angptl4*^fl/fl^ mice (**Figure 8B**). Again, *Angptl4*^AdipoKO^ male mice fed a NCD had increased uptake of radiolabeled fat into white adipose depots compared to floxed controls, but on HFD those differences largely disappeared (**Figure 8C**). No genotype-specific differences in tissue radiolabel uptake were observed in *Angptl4*^LivKO^ mice (**Figure 8D**).

**Figure 8.**
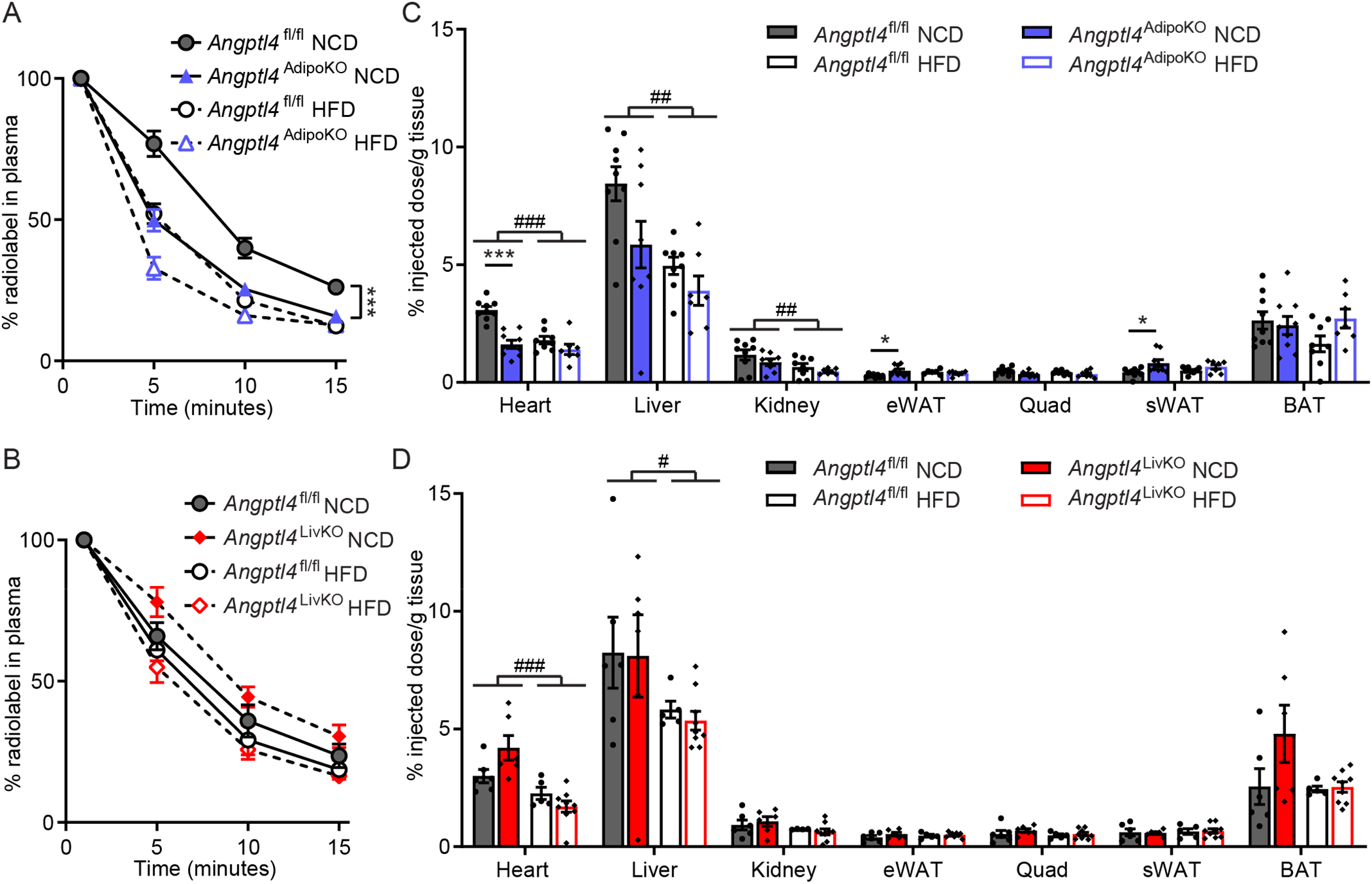
Chylomicron clearance and uptake and lipase activity in *Angptl4*^AdipoKO^ and *Angptl4*^LivKO^ mice after chronic high-fat feeding. At the conclusion of 6 months of either normal chow diet (NCD) or high fat diet (HFD) *Angptl4*^AdipoKO^ (A,C) and *Angptl4*^LivKO^ (B,D) mice (n=5-9/group) were fasted (6 h) and injected intravenously with ^3^H-triglyceride containing chylomicrons. **A and B)** Clearance of radiolabel from the plasma 1, 5, 10, and 15 minutes after injection. Points represent percentage of radiolabel remaining in the plasma at the indicated time points compared to the 1-minute time point (mean±SEM). ***p<0.001 by repeated measures ANOVA. **C and D)** Uptake of radiolabel (% injected dose/g tissue) into the indicated tissues after 15 min (mean±SEM). #p<0.05, ##p<0.01, ###p<0.001 for dietary differences by two-way ANOVA. *p<0.05, ***p<0.001 for individual genotype-specific differences by multiple comparison after two-way ANOVA.

To understand why adipose tissue deficiency no longer resulted in improved adipose clearance of triglycerides after a 6-month high-fat diet, we next measured tissue specific LPL activity. After 6 months of high-fat feeding, LPL activity was still greater in the adipose depots of *Angptl4*^AdipoKO^ mice compared to those of *Angptl4*^fl/fl^ mice (**Figure 9A**). These data suggest that during the setting of obesity increased LPL activity in the adipose tissue of *Angptl4*^AdipoKO^ mice no longer leads to increased adipose triglyceride uptake. This may be because the capacity of white adipose tissues to store triglycerides has become the limiting factor for adipose storage rather than the rate of LPL lipolysis. There were no genotype specific differences in LPL activity in the *Angptl4*^LivKO^ mice compared to floxed control mice (**Figure 9B**). Similar to what was observed after 12 weeks on HFD, there were no genotype-specific differences in total lipase, hepatic lipase, or LPL activity in the livers of either *Angptl4*^AdipoKO^ or *Angptl4*^LivKO^ mice (**Figure 9C-F**). We also measured tissue gene expression of *Lpl* and found no genotype-specific differences (**Supplemental Figure 10**).

**Figure 9.**
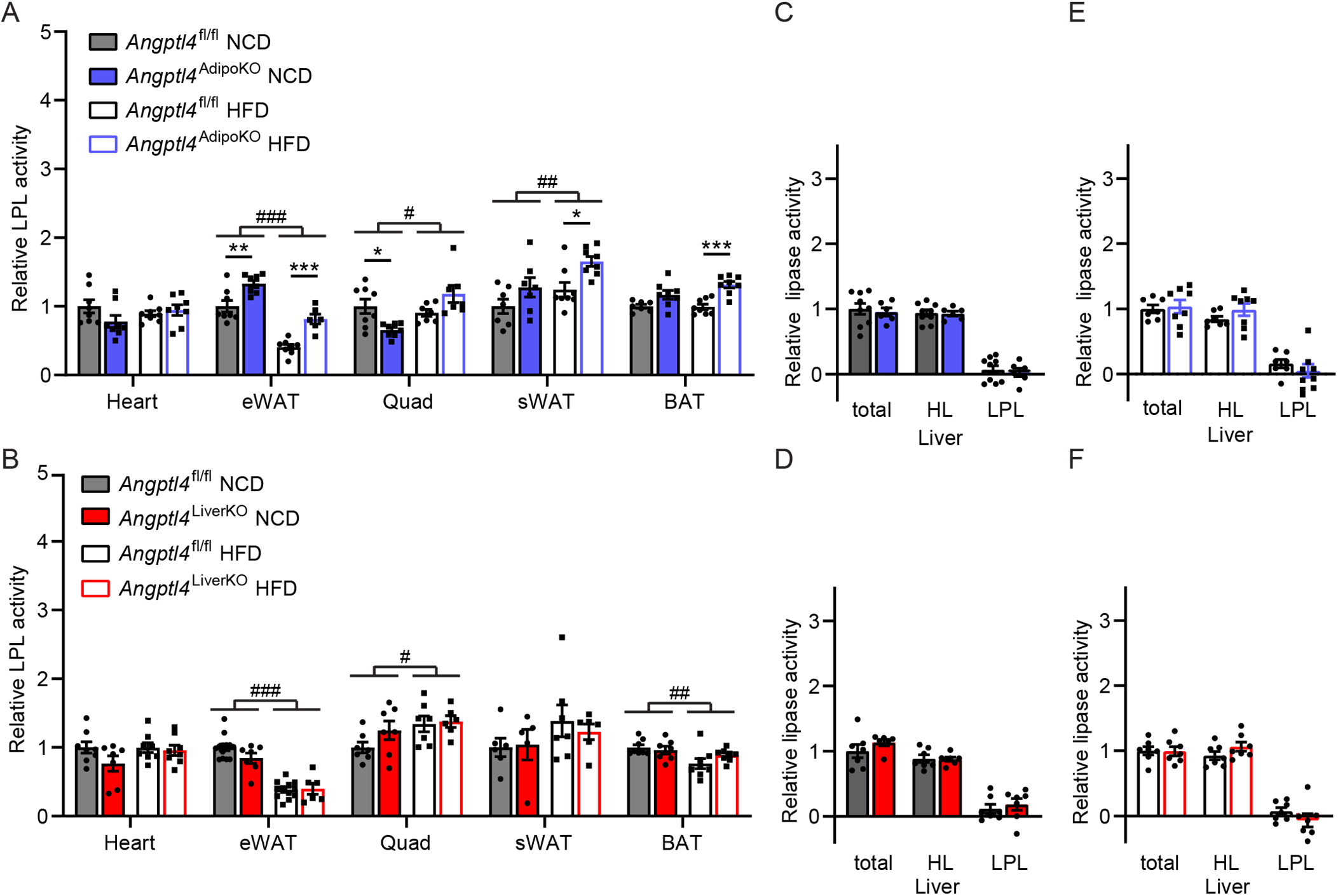
Lipase activity in *Angptl4*^AdipoKO^ and *Angptl4*^LivKO^ mice after chronic high-fat feeding. **A and B)** Heart, epididymal adipose tissue (eWAT), Quadricep muscle (Quad), subcutaneous adipose tissue (sWAT), and brown adipose (BAT) tissue) from fasted (6 h) male *Angptl4*^AdipoKO^ (E) and *Angptl4*^LivKO^ (F) mice (n=4-13/group) were harvested and lipase activity was measured. **C–F)** Liver was harvested from fasted (6 h) male *Angptl4*^AdipoKO^ (C, NCD groups and E, HFD groups) and *Angptl4*^LivKO^ (D, NCD groups and G, HFD groups) mice (n=6-8/group). Lipase activity was measured in the presence or absence of 1M NaCl to distinguish between hepatic and lipoprotein lipase. Bars show relative lipase activity in each tissue normalized to *Angptl4*^fl/fl^ (mean±SEM). #p<0.05, ##p<0.01, ###p<0.001 for dietary differences by two-way ANOVA. *p<0.05, **p<0.01 ***p<0.001 for individual genotype-specific differences by multiple comparison after two-way ANOVA.

Finally, we asked if the disappearance of improved adipose triglyceride clearance after 6 months on HFD affected the improved glucose tolerance we had observed in *Angptl4*^AdipoKO^ mice after 12 weeks on HFD (see Figure 6). Indeed, we found that *Angptl4*^AdipoKO^ mice no longer had improved glucose tolerance after 6 months on HFD (**Figure 10A**). And while *Angptl4*^AdipoKO^ mice remained more insulin sensitive than *Angptl4*^fl/fl^ mice after 6 months on HFD, the improvement was greatly reduced compared to that observed after 12 weeks of HFD (**Figure 10C**). Interestingly, despite no evidence of altered triglyceride partitioning, *Angptl4*^LivKO^ mice displayed improved glucose tolerance compared to littermate *Angptl4*^fl/fl^ controls after 6 months of high-fat feeding (**Figure 10B**). Insulin sensitivity was also slightly, but not statistically significantly improved (**Figure 10D**). As before, we measured tissue expression (liver, eWAT, sWAT, and BAT) of inflammatory markers *Ccl2, Cd68* and *Tnfα*. Once again, there was an increase in inflammation in HFD-fed mice in tested tissues, but no major genotype specific differences were observed (**Supplemental Figure 11**).

**Figure 10.**
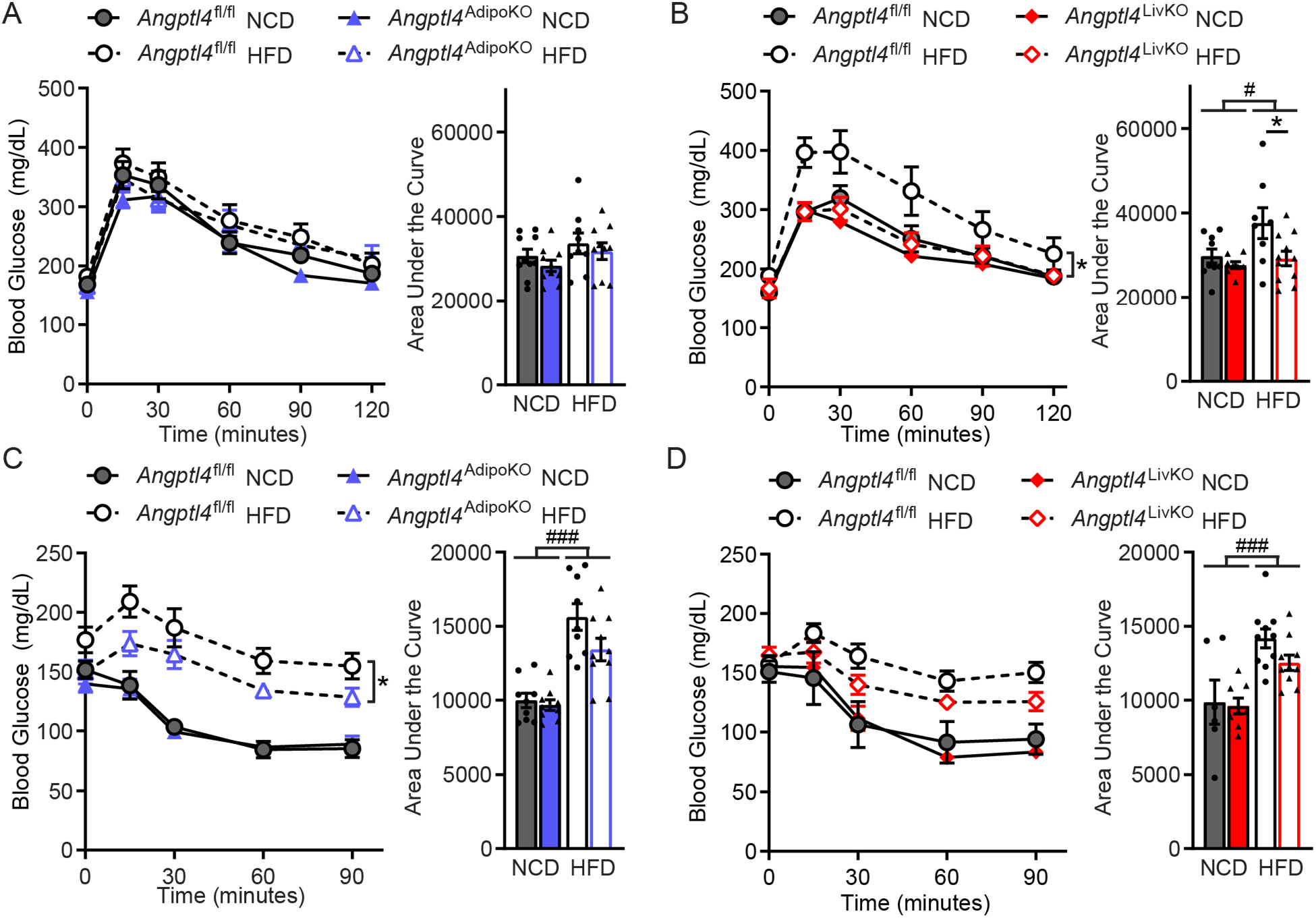
Glucose tolerance and insulin sensitivity of *Angptl4*^AdipoKO^ and *Angptl4*^LivKO^ mice after chronic high fat feeding. **A and B)** Glucose tolerance tests were performed on fasted (6 h) male *Angptl4*^AdipoKO^ (A) and *Angptl4*^LivKO^ (B) mice after 24 weeks of either a normal chow diet (NCD) or high fat diet (HFD). Mice were injected with glucose (1 g/kg) and blood glucose concentrations were measured over 2 h. Points represent glucose levels (mean±SEM; n=6-10) at each respective time point. Bar graphs represent area under the curve (mean±SEM) for all time points. *p<0.05 by repeated measures ANOVA. **C and D)** Insulin tolerance tests were performed on fasted (4 h) male *Angptl4*^*AdipoKO*^ (C) and *Angptl4*^*LivKO*^ (D) mice after 25 weeks of either a normal chow diet (NCD) or high fat diet (HFD). Mice were injected with 0.75 U/ml of human insulin (Humalin-R) and blood glucose concentrations were measured over 90 min. Points represent glucose levels (mean±SEM; n=5-9) at each respective time point. Bar graphs represent area under the curve (mean±SEM) for all time points. #p<0.05, ###p<0.001 for dietary differences by two-way ANOVA. *p<0.05, for individual genotype-specific differences by multiple comparison after two-way ANOVA (Tukey correction).

During the completion of this work another study investigating the role of liver ANGPTL4 has appeared in preprint. Singh *et al.* used the KOMP flox allele to generate a liver-ANGPTL4 deficient mouse model. Contrary to our observations, they observed lower levels of fasting plasma TG levels in their liver-deficient *Angptl4* mice fed either a normal chow or a Western diet for 16 weeks (21). Because Singh *et al.* measured plasma TG levels after an overnight fast and we measured TG levels after a 6 h fast, we asked if fasting our *Angptl4*^LivKO^ mice for 18 h would uncover a change in plasma TG levels. Indeed, after prolonged fasting our *Angptl4*^LivKO^ mice had decreased plasma TG levels compared to *Angptl4*^fl/fl^ mice, but when plasma TG levels were measured in the same mice after a 6 h fast no difference were observed, consistent with our previous observations (**Figure 11A**). Plasma TG levels following an 18 h fast were no longer different between *Angptl4*^LivKO^ and *Angptl4*^fl/fl^ mice fed HFD for 6 months, but lower plasma TG levels were still observed in age-matched mice fed a NCD (**Figure 11B**). We also performed a TG clearance and uptake assay in 8-12 week old male *Angptl4*^LivKO^ and *Angptl4*^fl/fl^ mice after an overnight fast. Although there was a trend towards increased radiolabeled TG clearance in the *Angptl4*^LivKO^ mice compared to *Angptl4*^fl/fl^ mice (**Figure 11C**), no differences in tissue TG uptake were observed (**Figure 11D**).

**Figure 11.**
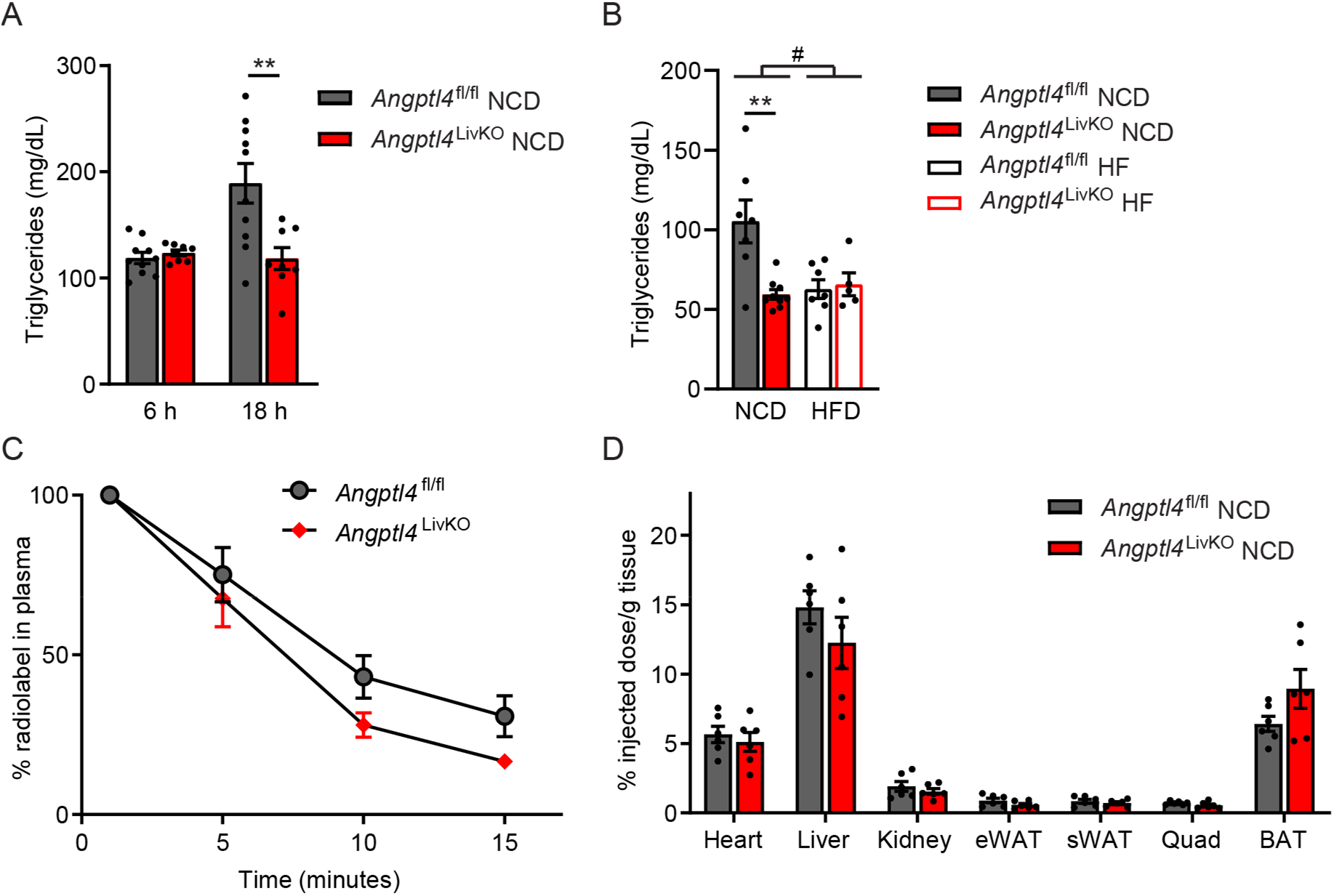
Triglyceride levels and partitioning in *Angptl4*^LivKO^ mice after overnight fast. **A)** Plasma triglyceride levels in fasted (6 h and 18 h) 8 to 12–week-old male *Angptl4*^fl/fl^ and *Angptl4*^LivKO^ mice (mean±SEM; n=8-10). **B)** Plasma triglyceride levels in fasted (18 h) *Angptl4*^fl/fl^ and *Angptl4*^LivKO^ mice after 26 weeks of either a normal chow diet (NCD) or high fat diet (HFD) (mean±SEM; n=7-9). **C)** Clearance of radiolabel from the plasma 1, 5, 10, and 15 minutes after intravenous injection of ^3^H-labeled chylomicrons in 8-12 week old male *Angptl4*^fl/fl^ and *Angptl4*^LivKO^ mice fasted for 18 h (mean±SEM; n=7-9). Points represent percentage of radiolabel remaining in the plasma at the indicated time points compared to the 1-minute time point (mean±SEM). **D)** Uptake of radiolabel (% injected dose/g tissue) into the indicated tissues after 15 min (mean±SEM). #p<0.05 for dietary differences by two-way ANOVA. **p<0.01 for individual genotype-specific differences by multiple comparison after two-way ANOVA.

## DISCUSSION

Proteins made and secreted by the liver (hepatokines) and adipose (adipokines) tissues play a vital role in lipid and glucose homeostasis during obesity. ANGPTL4 is highly expressed by both the adipose and liver in mice and humans (3, 22, 23). In this study we investigated the triglyceride phenotypes of adipocyte and hepatocyte-specific ANGPTL4 knockout mice. We also investigated how adipocyte or hepatocyte-specific loss of ANGPTL4 affected body weight, glucose tolerance, and insulin resistance in the setting of either 3 or 6 months of high-fat diet feeding and obesity. We found that adipocyte-, but not hepatocyte-specific loss of ANGPTL4 resulted in lower plasma triglycerides and increased uptake of triglyceride-fatty acids into adipose tissue. We also found that for mice fed a high-fat diet, adipocyte-specific ANGPTL4 deletion initially resulted in improved glucose tolerance and insulin sensitivity, but over time this improvement disappeared. Conversely, hepatocyte-specific knockout mice initially showed no difference in glucose tolerance and insulin sensitivity, but after 6 months on HFD, these mice showed signs of improvement in both.

The role of ANGPTL4 in regulating triglyceride metabolism through LPL inhibition has been well established. Human carriers of the E40K mutation in ANGPTL4 have lower triglycerides and a lower risk of coronary artery disease (6) and dyslipidemia during obesity (9). In mice, genetic loss of ANGPTL4 leads to decreased fasting plasma TG levels and increased uptake of triglycerides into adipose tissue (3, 14, 15, 19). The specific increase in adipose tissue LPL activity and TG uptake in whole body ANGPTL4 knockout mice strongly supported the idea that adipose derived ANGPTL4 was responsible for the triglyceride phenotypes in whole-body ANGPTL4 knockout mice. Here, adipocyte-specific loss of ANGPTL4 in mice fed a chow diet phenocopies the triglyceride phenotypes observed in whole-body *Angptl4*^−/−^ mice. *Angptl4*^AdipoKO^ mice had lower plasma triglyceride levels, increased uptake of triglycerides into adipose tissue, and increased adipose LPL activity. In contrast, hepatocyte-specific knockout of ANGPTL4 resulted in few discernable triglyceride phenotypes, at least under the conditions tested in this study. This was true in both male and female mice. The triglyceride phenotypes observed in *Angptl4*^AdipoKO^ mice were consistent with those recently reported by Aryal *et al.* using an independently derived adipocyte-specific ANGPTL4 knockout mouse (14). It is important to note that unlike the flox allele used by Aryal *et al,* which is predicted, after recombination, to yield a truncated ANGPTL4 containing the first 1-186 amino acid residues and may retain hypomorphic function (23), our allele did not appear to yield any protein. Both the current study and the Aryal *et al*. study support the hypothesis that it is primarily loss of adipose ANGPTL4 that is responsible for the alterations in plasma triglyceride levels and triglyceride partitioning observed in *Angptl4*^−/−^ mice.

Loss of adipocyte ANGPTL4 in the context of HFD had an interesting effect on LPL activity and adipose triglyceride uptake. On a chow diet, the increase in adipose LPL activity in *Angptl4*^*AdipoKO*^ mice led to a corresponding increase in adipose-tissue uptake of triglycerides. This connection between increased LPL activity and increased triglyceride uptake in adipose was lost in mice fed a HFD. Adipose LPL activity remained higher in the adipose tissue of *Angptl4*^*AdipoKO*^ mice fed HFD compared to floxed control mice, but the enhanced adipose-tissue uptake observed on chow diet was largely lost. These data suggest that on a chronic high-fat diet, LPL activity is no longer rate-limiting for the entry of triglyceride-derived fatty acids into adipose. We postulate that this is likely due to the adipose dysfunction and chronic inflammation that accompany obesity and high-fat feeding (20, 24).

High-fat feeding experiments performed by Janssen *et al*. (25) suggested that ANGPTL4 deficiency not only impacts triglyceride metabolism, but also glucose metabolism. Their work demonstrated that *Angptl4*^−/−^ mice fed a diet rich in unsaturated fatty acids, gained more weight than wildtype mice, but had lower blood glucose levels and improved glucose tolerance. Indeed, human studies found that carriers of the inactivating E40K mutation in *Angptl4* had lower fasting glucose levels than non-diabetic participants and had a lower odds ratio (0.89) for developing type 2 diabetes (26). One hypothesis is that improved glucose homeostasis is due to relief of LPL inhibition and improved triglyceride clearance to adipose (25). We did find that *Angptl4*^AdipoKO^ mice had increased clearance of triglycerides to adipose and that after 12 weeks on HFD, these mice have improved glucose tolerance and are protected from insulin resistance. However, the improved glucose tolerance and protection from insulin resistance were mostly lost after long term HFD feeding. These observations are consistent with recent reports where independent strains of total adipocyte-specific or brown adipocyte–specific ANGPTL4 deficient mice demonstrated improved glucose tolerance after short-term HFD feeding (4 weeks), but lost these improvements after 20 weeks of HFD feeding (14, 18). As discussed above, we found that chronically HFD fed *Angptl4*^AdipoKO^ mice did not have increased uptake of triglycerides into adipose tissue. We suspect that increasing TG partitioning to adipose improves glucose homeostasis, but that when the increased TG uptake to adipose is lost, improved glucose tolerance is also lost. Therapeutically, these findings imply that targeting ANGPTL4 to improve glucose tolerance may not be effective over the long-term, at least not in the situation of continued overconsumption of calories.

A major question left unanswered in our study is the role of ANGPTL4 produced by hepatocytes. Compared to littermate controls, after a 6 h fast we observed no differences in plasma triglyceride levels or triglyceride uptake levels in *Angptl4*^LivKO^ mice fed either HFD or normal chow. We also observed no differences in LPL activity, hepatic lipase activity, liver size, or liver triglycerides in these mice. After a prolonged (18 h) fast, however, we did observe lower triglyceride levels in *Angptl4*^LivKO^ mice. The physiological relevance of this observation is unclear. Even after an 18 h fast, we observed no increase in TG uptake into metabolically active tissues. Fasting overnight or longer in mice induces a catabolic, starvation-like state and is not equivalent to an overnight fast in humans (27, 28), thus our preference for measuring plasma lipid levels after a 6 h fast. Fasting alters enzyme activities and expression of many genes important to lipid metabolism (29), and it has been shown that liver weights, liver triglyceride content, liver enzyme activities, and expression of genes related to lipogenesis and lipolysis can differ significantly depending on the length of fasting (30). We have previously shown that *Angptl4* expression in mouse tissues, including the liver, is induced within 2 h of fasting (15). The fact that we did not observe changes in plasma triglyceride levels until much later in a fast suggests that perhaps liver ANGPTL4 is not acting directly on triglyceride clearance to lower plasma triglyceride levels.

Interestingly, we did find some evidence that liver ANGPTL4 does affect overall energy homeostasis. After 6 months of HFD feeding, glucose tolerance was improved and there was a trend towards improved insulin sensitivity. These improvements occurred in the absence of differences in fasting plasma TG levels or alterations in tissue TG uptake. Moreover, metabolic caging data indicated that *Angptl4*^LivKO^ mice have increased energy expenditure. The mechanisms behind these improvements are unclear. Given that the improved glucose tolerance emerged only in mice fed a chronic HFD, it would be interesting to determine if an even longer period of HFD feeding would increase the difference in glucose tolerance between *Angptl4*^LivKO^ mice and controls, and if the increased energy expenditure we observed in *Angptl4*^LivKO^ mice would persist at older ages.

The current study is not without limitations. We chose a 60% (kCal) high fat diet, a diet that is widely used experimentally to induce an obese and insulin resistant phenotype. Other diets may more closely reflect the typical American diet, such as the “Western diet” comprised of 42% (kCal) high fat, high sucrose and high cholesterol. Alternative diets that induce different pathological states might illuminate tissue-specific roles for ANGPTL4 that were missed in our study. Another limitation to our study is that we studied adipocyte- or hepatocyte-specific *Angptl4* knockout separately, but not in combination. Therefore, we would have missed any synergistic effects of ANGPTL4-deficiency in both tissues. Generating mice that are deficient both in hepatocyte and adipocyte ANGPTL4 might allow for a greater interrogation of ANGPTL4’s role in obesity, metabolic syndrome, and insulin resistance while likely still circumventing the HFD-induced lethal phenotype observed in whole body *Angptl4*^−/−^ mice. ANGPTL4 is expressed, although at lower levels, in skeletal muscle and heart (31) and could have important actions during exercise (32) or situations of lipotoxicity (33). ANGPTL4 expressed in the intestine has also been implicated in the effects of the intestinal microbiota on metabolism (25). Generating additional tissue-specific *Angptl4* knockout mice could identify the roles of ANGPTL4 in these and other tissues.

Our studies highlight the importance of studying age and diet duration going forward. HFD studies in mice are generally performed in mice that are less than 1 year old and who have been fed a high-fat diet for 4 to 16 weeks. However, both Aryal et al. (14) and our current study clearly showed that improved glucose tolerance in adipocyte-specific ANGPTL4 knockout mice disappeared with chronic high-fat feeding. Moreover, we found that improvements in glucose homeostasis in our *Angptl4*^LivKO^ mice only emerged *after* long-term HFD feeding. Given that metabolic disease is most prevalent in older and obese populations, these observations highlight the need and importance of performing long-term feeding studies in aged mice if we are to fully understand the roles for different proteins in dyslipidemia and metabolic disease.

## METHODS

### Mice

Mice with floxed alleles of the *Angptl4* gene were generated by the University of Iowa genome editing facility using CRISPR/Cas9. B6SJLF1/J mice were purchased from Jackson Labs (100012; Bar Harbor, ME). Male mice older than 8 weeks were bred with 3 to 5–week-old super-ovulated females to produce zygotes for microinjection. Female ICR (Envigo; Hsc:ICR(CD-1)) mice were used as recipients for embryo transfer. Chemically modified CRISPR-Cas9 crRNAs and tracrRNAs were purchased from IDT (Alt-R® CRISPR-Cas9 crRNA; Alt-R® CRISPR-Cas9 tracrRNA (Cat# 1072532)). crRNA sequences were CCCTTTCACAGTCTGCTCTG and TGTGTCTAGTCTAGGAGCCG. The crRNAs and tracrRNA were suspended in T10E0.1 buffer (10 mM Tris pH 8, 0.1 mM EDTA) and combined to 1 μg/μl (~29.5 μM) final concentration in a 1:2 (μg:μg) ratio. The RNAs were heated at 98°C for 2 min and allowed to cool slowly to 20°C in a thermal cycler. The annealed cr:tracrRNAs were aliquoted to single-use tubes and stored at −80°C. Cas9 nuclease was also purchased from IDT (Alt-R® S.p. HiFi Cas9 Nuclease). Individual cr:tracr:Cas9 ribonucleoprotein complexes were made by combining Cas9 protein and cr:tracrRNA in T10E0.1 (final concentrations: 100 ng/ul (~0.6 μM) Cas9 protein and 100 ng/μl (~2.9 μM) cr:tracrRNA). The Cas9 protein and annealed RNAs were incubated at 37°C for 10 minutes. The two RNP complexes were mixed resulting in a final concentration of 100 ng/μl (~0.6 μM) Cas9 protein and 50 ng/μl (~1.5 μM) each cr:tracrRNA. Long single stranded DNA was prepared by digesting the *Angptl4* target vector (524 bp 5’ arm of homology; 911 bp floxed region; 636 bp 3’ arm of homology) with the nicking enzyme Nb.Bsm1 (New England Biolabs) and purification from an alkaline-agarose gel. The final concentrations in the micro-injection mix were 20 ng/μl each cr:tracr RNA (~0.6 μM), 40 ng/μl Cas9 protein (~0.3 μM), and 10 ng/μl long ssDNA. Pronuclear-stage embryos were collected using methods described in (34). Embryos were collected in KSOM media (Millipore; MR101D) and washed 3 times to remove cumulous cells. Microinjection into the male pronucleus was performed and embryos incubated overnight. Two cell zygotes were implanted into pseudo-pregnant ICR females.

All pups were tested for the appropriate insertion of 5’ and 3’ LoxP sites by PCR amplification of the genomic region and subsequent sequencing. The above procedures resulted in the successful generation of a single male mouse with both 5’ and 3’ LoxP sites. This mouse was bred to a C57Bl/6 female mouse and the heterozygous progeny were interbred to generate homozygous floxed mice. During breeding, the presence of the LoxP Sites was confirmed by genotyping PCR using oligonucleotides TAGGCGCATCTACTAGGACTC (forward) and AGATATGCAAGGCTAGTGAAGAC (reverse) to detect the 5’ LoxP site and oligonucleotides CCTCCAACATCTCTTGATGTAAC (forward) and ATATGTGTATGTGACTGGATGG (reverse) to detect the 3’ LoxP site. Adipocyte-specific knockout mice (*Angptl4*^AdipoKO^) were generated by breeding *Angptl4*^fl/fl^ mice with C57/BL6 transgenic mice containing the adiponectin promoter-driven Cre recombinase (kindly provided by Dr. Matthew Potthoff, Jackson stock 010803 (35)). Hepatocyte-specific knockout mice (*Angptl4*^LivKO^) were generated by breeding *Angptl4*^fl/fl^ mice with C57/BL6 transgenic mice containing the albumin promoter-driven Cre recombinase (kindly provided by Dr. Eric Taylor, Jackson stock 003574 (36)). The presence of adiponectin-Cre was verified by genotyping PCR using GGATGTGCCATGTGAGTCTG (forward) and ACGGACAGAAGCATTTTCCA (reverse) as primers. Primers used to confirm albumin-Cre by genotyping PCR were TGCAAACATCACATGCACAC (forward to detect wildtype), GAAGCAGAAGCTTAGGAAGATGG (forward to detect albumin Cre), and TTGGCCCCTTACCATAACTG (common reverse). In all studies Cre-negative *Angptl4*^fl/fl^ littermates were used as controls.

All animal procedures were approved by the Institutional Animal Care and Use Committee at the University of Iowa and were carried out in accordance with the National Institute of Health *Guide for Care and Use of Laboratory Animals*. All animals were maintained in a climate-controlled environment at 25°C with a 12:12 hour light-dark cycle. Water was provided *ad libitum*. During non-fasting conditions mice were given standard mouse chow (NIH 7951) until 8 weeks of age when high-fat diet studies began. At 8 weeks of age mice were assigned to either remain on standard mouse chow (NIH 7951) or given high-fat diet (60% by kCal; Research Diets, 12592). Mice were fed their respective diets for either 12 weeks or 6 months.

### Production of *Angptl4* and LPL conditioned media

A construct expressing full-length mouse ANGPTL4 (pXC4) was generated from full-length mouse ANGPTL4 cDNA (MMM1013-202763591). A V5 tag was appended to the C terminus of the open reading frame using Phusion site-directed mutagenesis (New England Biolabs) to generate a V5 tagged version of mouse ANGPTL4 (pHS5). To generate a mouse ANGPTL4 construct mimicking the predicted protein product of our *Angptl4* flox allele after Cre-mediated recombination (pKMS22) we deleted residues 111-187 (Exons 2 and 3) from full-length mouse ANGPTL4 by site-directed mutagenesis. A V5 tag was appended to the new C terminus of the altered open reading frame using site directed mutagenesis (pKMS25). We also generated a mouse ANGPTL4 construct to mimic the predicted protein product of a flox allele generated by the EUCOMM/KOMP consortium and used by several previous studies (14, 17, 18). This independently generated flox *Angptl4* allele uses a knock-out first strategy and has LoxP sites inserted into the *Angptl4* gene flanking Exons 4,5, and 6. To generate a construct mimicking the predicted protein after Cre-mediated recombination (pXC32) we deleted the residues coded by exons 4 through 6 from full-length mouse ANGPTL4 by site-directed mutagenesis. A V5 tag was appended to the C terminus also using site directed mutagenesis (pXC34). To produce conditioned media containing either V5-tagged mANGPTL4, V5-tagged Flox mANGPTL4 or V5-tagged KOMP allele mANGPTL4, 293T cells were transiently transfected with the respective constructs using the transfection agent PEI. 24 h post-transfection the media was switched to serum-free DMEM containing 1× protease inhibitors and grown for 48 h. Mock transfected 293T cells were used as a control. The media was then collected for use in Western blot analysis and LPL activity assays. FLAG-tagged human LPL was concentrated from the medium of a Chinese hamster ovary cell line (CHO-K1) stably expressing FLAG-tagged human LPL as previously described (37, 38). The presence of LPL in the conditioned media was confirmed through Western blotting using a mouse antibody against the FLAG tag (1:5000; Sigma). LPL activity was assessed through an LPL activity assay (see below).

### Western Blot

Lysate and media samples collected from 293T cells transfected with V5-tagged mANGPTL4, V5-tagged Flox mANGPTL4, or V5-tagged KOMP allele mANGPTL4 constructs were size fractionated on 12% SDS-polyacrylamide gels and then transferred to nitrocellulose membranes. Membranes were blocked with casein before primary antibodies were added (1:3000 dilution of mouse monoclonal antibody against the V5 tag (ThermoFisher MA5-15253)) in a casein buffer containing Tween. Membranes were rocked overnight at 4°C. Following incubation, the primary antibody was removed and membranes washed 3 times with PBS containing 0.1% Tween. Membranes were then incubated with anti-mouse Dylight800-labeled secondary antibody (Invitrogen) diluted 1:5000 in casein buffer. After washing with PBS containing Tween, antibody binding was detected using an Odyssey Infrared Scanner (LI-COR).

### Analysis of Fasting Plasma Parameters

Fasting plasma parameters were measured in fasted (6 h) *Angptl4*^fl/fl^, *Angptl4*^LivKO^, and *Angptl4*^AdipoKO^ mice. Blood was collected into EDTA-coated capillary tubes following a tail-nick and centrifuged to collect plasma (1,500×g, 15 minutes, 4°C). Plasma TG, NEFA, and glucose measurements were taken at 8 weeks of age before beginning diet and 12 weeks or 6 months following the start of diet studies. For plasma TG analysis, plasma (2 μl, technical duplicates using the same aliquot of plasma per individual mouse were used) was combined with Infinity^TM-^ Triglyceride Reagent (200 ul, Thermo Scientific TR22421) in a 96 well plate. Samples were incubated for 10 min at 37°C, gently tapped to ensure proper mixing of reagent with sample, and absorbance was read at 500 nm and 660 nm. Triglyceride concentrations were determined using a standard curve prepared from Triolein standards (Nu-Chek Prep, Lot T-235-N13-Y). For plasma NEFA analysis, plasma (2 μl, technical duplicates) was combined with Color Reagent A (112.5 μl, Wako HR Series NEFA-HR 999-34691/995-34791) and incubated for 10 min at 37°C. Following incubation, Color Reagent B was added to samples (37.5 μl, Wako HR Series NEFA-HR 991-34891/993-35191) and incubated for 5 min at 37°C. Absorbance was read at 560 nm and 670 nm. Non-esterified fatty acid concentrations were determined using a standard curve prepared from Oleic Acid standard (TCI, O0180, Lot Z8RVM). Fasting plasma glucose concentration was measured once per mouse immediately following a tail nick using a glucometer (One Touch Ultra). Plasma amyloid A was measured using an ELISA kit (Crystal Chem #80659) following manufacturer’s protocol. Plasma activity of aspartate aminotransferase (AST, Sigma-Aldrich, MAK055) and alanine aminotransferase (ALT, Sigma-Aldrich, MAK052) were measured according to manufacturer’s protocols.

### RNA Extraction and qPCR Analysis

Mouse tissues (heart, liver, gonadal and epididymal white adipose tissue, quadriceps muscle, subcutaneous white adipose tissue and brown adipose tissue) were harvested following a 6 h fast. Tissue was snap frozen in liquid nitrogen and stored at −80°C until processed. Tissues were pulverized before homogenization. Total RNA was extracted using Trizol (Invitrogen) according to manufacturer’s instructions. 2 μg of RNA was used to prepare cDNA with the High Capacity cDNA Reverse Transcription kit (Applied Biosystems, 4368813) according to manufacturer’s instructions. Prepared cDNA was used for qPCR analysis using primers for *Angptl4* (forward-TTTGCAGACTCAGCTCAAGG; reverse-TCCATTGTCTAGGTGCGTGG), *CycloA* (forward-TGGCAAGACCAGCAAGAA; reverse-CTCCTGAGCTACAGAAGGAATG), *Lpl* (forward-CTGGTCTTAACCGGCCCAAT; reverse-TGCACATAGCCAGAAGGGTG), *U36B4* (forward-CGTCCTCGTTGGAGTGACA; reverse-CGGTGCGTCAGGGATTG), *Ccl2* (forward-CCCAATGAGTAGGCTGGAGA; reverse-TCTGGACCCATTCCTTCTTG), *Cd68* (forward-GGGGCTCTTGGGAACTACAC; reverse-GTACCGTCACAACCTCCCTG), and *Tnfα* (forward-CCCTCACACTCAGATCATCTTCT; reverse-GCTACGACGTGGGCTACAG). Primers for Supplemental Figure 1 are in listed in Table 1 of Supplemental Data. Prepared diluted cDNA, primers, and SYBR Green ER qPCR Supermix reagent (Invitrogen, 11762100) were combined, and PCR was performed on the QuantStudio 6 Flex system (3 technical replicates per mouse, Applied Biosystems, Iowa Institute of Human Genetics). Gene expression was calculated using the ΔΔct method (39) using either *CycloA* or *U36B4* as the reference gene.

### Preparation of ^3^H-Labeled Chylomicrons

^3^H-labeled chylomicrons were prepared as previously described (15). Briefly, glycosylphosphatidylinositol-anchored high-density lipoprotein binding protein 1 (GPIHBP1) knock out mice were orally gavaged with 100 μCi of [9,10-3H(N)]-Triolein (Perkin Elmer, NET431001MC) suspended in olive oil. After 4 h, blood was collected via cardiac puncture and placed into a tube containing 0.5 M EDTA and placed on ice. The blood was then centrifuged at 1,500×g for 15 min and plasma was collected. The plasma was then centrifuged at 424,000×g twice for 2 hours at 10°C. The chylomicrons were collected from the upper layer of the resulting supernatant, resuspended in sterile saline and stored at 4°C. Radioactivity was measured using a Beckman-Coulter Liquid Scintillation Counter.

### Triglyceride Uptake Assay

TG clearance and uptake assays were performed in mice as previously described (15). Briefly, mice were fasted for 6 hours and then injected retro-orbitally with 100 μl of ^3^H-labeled chylomicron suspension. Blood samples were taken 1, 5, 10, and 15 minutes after injection. For each time point, 10 μl of blood was assayed for radioactivity in BioSafe II scintillation fluid using a Beckman-Coulter Scintillation Counter. Following the final timepoint mice were anesthetized with isoflurane and perfused with 20 mL of cold 0.5% Tyloxapol in PBS solution. Tissues (heart, liver, kidney, gonadal or epidydimal white adipose tissue, quadriceps, subcutaneous white adipose tissue, and brown adipose tissue) were harvested and weighed. 40-90 mg of each tissue was placed into a glass vial containing a 2:1 chloroform:methanol solution and stored overnight at 4°C. 1 mL of 2 M CaCl_2_ was added to each vial to separate the organic and aqueous phases. Samples were centrifuged for 10 min at 1,000 rpm and the upper aqueous layer was assayed for radioactivity in BioSafe II scintillation fluid using a Beckman-Coulter Scintillation Counter. The organic layer was allowed to evaporate overnight in an empty scintillation vial, then resuspended in scintillation fluid and counted. The counts per million (CPM) from the aqueous and organic fractions were combined to obtain the total uptake CPMs. To normalize radiolabel across mice, CPM values were normalized to the CPMs of the injected dose (measured by assaying 10% of the chylomicron suspension injected into the mouse).

### Liver Triglyceride Measurements

Liver triglyceride measurement was performed using a modified Folch extraction method (40). Liver tissues (approximately 100 mg) were homogenized in 500 μl of ice-cold PBS. 1.5 ml of a 2:1 (chloroform:methanol) mixture was added to each sample. Samples were vortexed briefly and incubated for 10-15 min. Samples were then centrifuged for 10 min at 2,050×g at 4°C. The lower organic phase was separated with a glass Pasteur pipette and placed into a 7 mL plastic vial (RPI 125509) and allowed to dry. The dried lipid was then dissolved with 500 μl of 2% Triton X-100 in chloroform and allowed to dry. The dried lipid was resuspended in 100 μl of molecular grade water. The sample was diluted 1 to 20 for triglyceride assay analysis. Analysis was performed using 2 μl of diluted lipid (technical duplicates) which was combined with Infinity^TM-^ Triglyceride Reagent (200 μl, Thermo Scientific TR22421) in a 96 well plate. The plate was incubated for 10 min at 37°C, gently tapped to ensure proper mixing of reagent with sample, and absorbance was read at 500 nm and 660 nm. Triglyceride concentrations were determined using a standard curve prepared from Triolein standard (Nu-Chek Prep, Lot T-235-N13-Y). Triglyceride levels were normalized to milligram of initial tissue weight.

### Lipoprotein Lipase Activity Assays

LPL activity in tissues was measured as previously described (15). Briefly, frozen tissue samples were crushed using a metal tissue pulverizer. The tissue was resuspended in LPL assay buffer (25 mM NH_4_Cl, 5 mM EDTA, 0.01% SDS, 45 U/mL heparin, 0.05% 3-(N,N-Dimethylmyristylammonio) propanesulfonate zwittergent detergent (Acros Organics, 427740050)) containing protease inhibitor (Mammalian ProteaseArrest-APExBIO K1008). The tissue was then vortexed and incubated on ice for 30 min with occasional further disruption with surgical scissors. The lysate was then clarified by centrifugation at 15,000×g for 15 min (4°C). Protein concentrations were equalized prior to assaying activity. Supernatants were combined with a buffer comprised of 0.6 M NaCl, 80 mM Tris-HCl pH 8, 6% fatty-acid free BSA, and 1% of the EnzChek lipase fluorescent substrate (Molecular Probes, E33955). Fluorescence was measured from technical duplicates of each lysate (30 min, 37°C) on a SpectraMax i3 plate reader (Molecular Devices). Relative lipase activity was determined following calculation of the linear slope of the curve and subtraction of background (assay buffer) slope readings. For liver lipase activity assays each lysate was treated with either vehicle (PBS) or NaCl (1M final concentration) to differentiate the roles of hepatic lipase (NaCl insensitive) from LPL (NaCl sensitive) before the addition of the assay buffer.

The LPL activity of LPL conditioned media treated with either V5-tagged mANGPTL4, V5-tagged Flox mANGPTL4, or V5-tagged KOMP allele mANGPTL4 constructs was measured as previously described (41). Briefly, ANGPTL4 containing conditioned media was combined with LPL conditioned media and samples were incubated at 37°C for 30 min. After incubation, 50 μl of each assay samples was combined with 25 μl of assay buffer (0.6 M NaCl, 80 mM Tris-HCl pH 8, 6% fatty-acid free BSA), following which 25 μl of substrate solution (1% of the EnzChek lipase fluorescent substrate (Molecular Probes, E33955) 0.05% 3-(N,N-dimethylmyristylammonio) propanesulfaonate Zwittergent detergent (Acros Organics) in 1% methanol) was then added to each sample. Fluorescence was measured from technical duplicates of each lysate (30 min, 37°C) on a SpectraMax i3 plate reader (Molecular Devices). Lipase activity was determined as stated above.

### Glucose and Insulin Tolerance Tests

Glucose tolerance tests (GTT) were performed after fasting mice for 6 h on week 10 on diet in the 12-week diet cohort or week 24 on diet in the 6-month diet cohort. Blood samples were collected before and 30, 60, 90, and 120 minutes following an intraperitoneal injection with glucose (12-week HFD study: 2g/kg for mice fed NCD, 1.3 g/kg for mice fed HFD; 6-month HFD study: 1g/kg for all mice). For mice fed a HFD for 12 weeks, blood was collected into an EDTA-coated capillary tube and stored on ice. Plasma was collected following centrifugation of sample tubes at 1,500×g for 20 minutes. Plasma glucose concentration was assessed using the Autokit Glucose kit (Wako-997-03001). 2.5 μl of plasma was combined with 200 μl of the Autokit glucose buffer solution and incubated at 37°C for 5 min. Absorbance was read at 505 nM and 600 nM. The 600 nM reading was subtracted from the 505 nM read. Glucose concentrations were determined using a standard curve prepared from glucose standards provided by the kit. For mice fed a HFD for 6 months, blood glucose readings were taken following a tail nick, before injection and 30, 60, 90, and 120 minutes following glucose injection using a glucometer (OneTouch Ultra).

Insulin tolerance tests (ITT) were performed following a 4 h fast in mice following 11 weeks on diet in the 12-week diet cohort and 25 weeks on diet in the 6-month diet cohort. Glucose readings were taken following a tail-nick bleed before and 15, 30, 60, and 90 minutes after an intraperitoneal injection with insulin (0.75U/kg, Humalin-R 100) using a glucometer (OneTouch Ultra).

### Metabolic Cage Studies

Mice were individually housed in Promethion cage systems from Sable Systems International at the University of Iowa Metabolic Phenotyping Core. Mice were allowed to acclimate to their housing for 48 hours. Metabolic measurements were then taken for 48 hours. Measurements were taken during the duration of two light cycles (6 a.m.–6 p.m.) and two dark cycles (6 p.m.–6 a.m.). Measurements were recorded as energy expenditure (kcal/hr), oxygen consumption (VO_2_ (ml/min)), carbon dioxide production (VCO_2_ (ml/min)) and respiratory quotient (RQ, VCO_2_/VO_2_).

### Body Composition

Body composition (fat and lean mass) was determined using nuclear magnetic resonance (NMR) in mice following 11 weeks on diet in the 12-week diet cohort and 25 weeks on diet in the 6-month diet cohort. Mice were weighed before being placed in a restraint tube without anesthesia and placed into either a Bruker LF50 (for mice under 50 grams) or a Bruker LF90 (for mice over 50 grams). Following NMR scanning mice were immediately returned to their cages. Body composition measurements were performed in the Fraternal Order of the Eagles Diabetes Research Center Metabolic Phenotyping Core.

### Statistics and Outlier Identification

Results are expressed as means ± SEM. Bar graphs also show individual values. Outlier identification was performed on all mouse datasets using ROUT analysis in GraphPad Prism. An unpaired Student’s *t*-test with Welch’s correction was used to determine statistical significance of samples with two groups (Figures 1 and 2). For groups of three or more statistical significance was determined by 2-way ANOVA followed by multiple comparison with Tukey correction. Repeated measured ANOVA was utilized for body weights, chylomicron clearance, GTT, and ITT assays. Statistical analysis was performed in GraphPad Prism.

## Author Contributions

BSJD and KMS conceived the study and designed the experiments. KMS conducted most of the experiments. SKS, EMC and KLDS conducted some experiments, acquired data and gave helpful suggestions. KMS and BSJD performed data analysis. KMS and BSJD wrote and edited the manuscript. All authors interpreted data, participated in manuscript review, and approved the final manuscript.

## Acknowledgements

This work was supported by grants from the National Institutes of Health (R01HL130146 [BSJD]). We thank the University of Iowa Genome Editing Facility for assistance in generating the ANGPTL4 floxed allele and the Fraternal Order of Eagles Diabetes Research Center Metabolic Phenotyping Core Laboratory for assistance is collecting metabolic caging data.

