## Supplemental data for "ANGPTL4 from adipose, but not liver, is responsible for regulating plasma triglyceride partitioning"

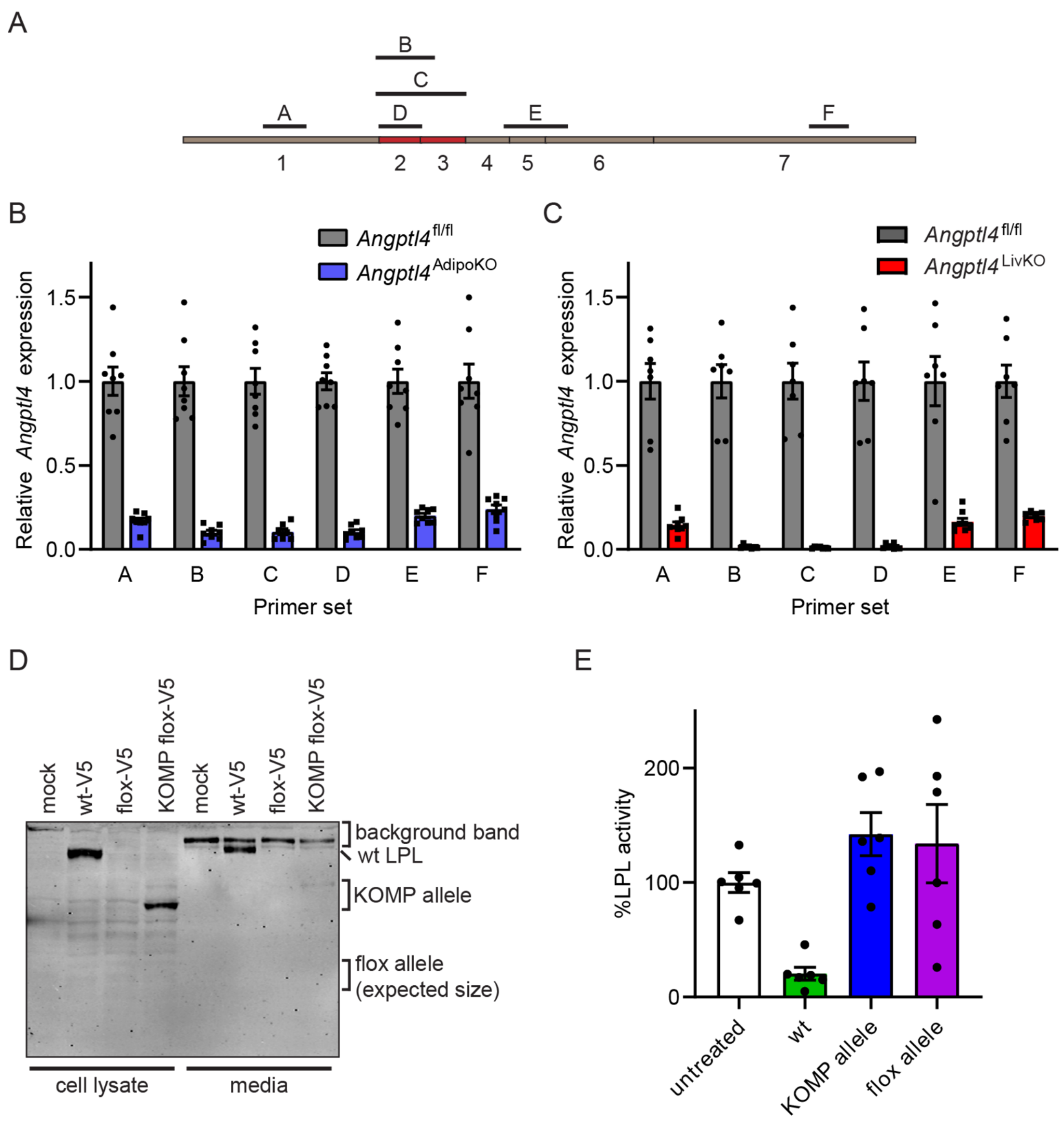


**Supplemental Figure 1: Characterization of *Angptl4* flox allele. A)** Schematic of the coding region of the ANGPTL4 gene showing the regions amplified by primers A-F. **B)** mRNA expression of *Angptl4* in brown adipose tissue of *Angptl4*$^{fl/fl}$ and *Angptl4*$^{AdipoKO}$ mice using primers A-F (mean±SEM; n=7-8). **C)** mRNA expression of *Angptl4* in liver tissue of *Angptl4*$^{fl/fl}$ and *Angptl4*$^{LivKO}$ mice using primers A-F (mean±SEM; n=7-8). **D)** Western blot of tissue lysate and conditioned media from 293T cells transfected with constructs encoding V5-tagged full length mouse ANGPTL4, V5-tagged Flox mouse ANGPTL4 or V5-tagged KOMP allele mouse ANGPTL4 probed with antibody against the V5 epitope. **E)** LPL activity of LPL treated with conditioned media from 293T cells transfected with constructs encoding V5-tagged full length mouse ANGPTL4, V5-tagged Flox mouse ANGPTL4, or V5-tagged KOMP allele mouse ANGPTL4 (means±SEM of three experiments with n=2 per group per experiment).

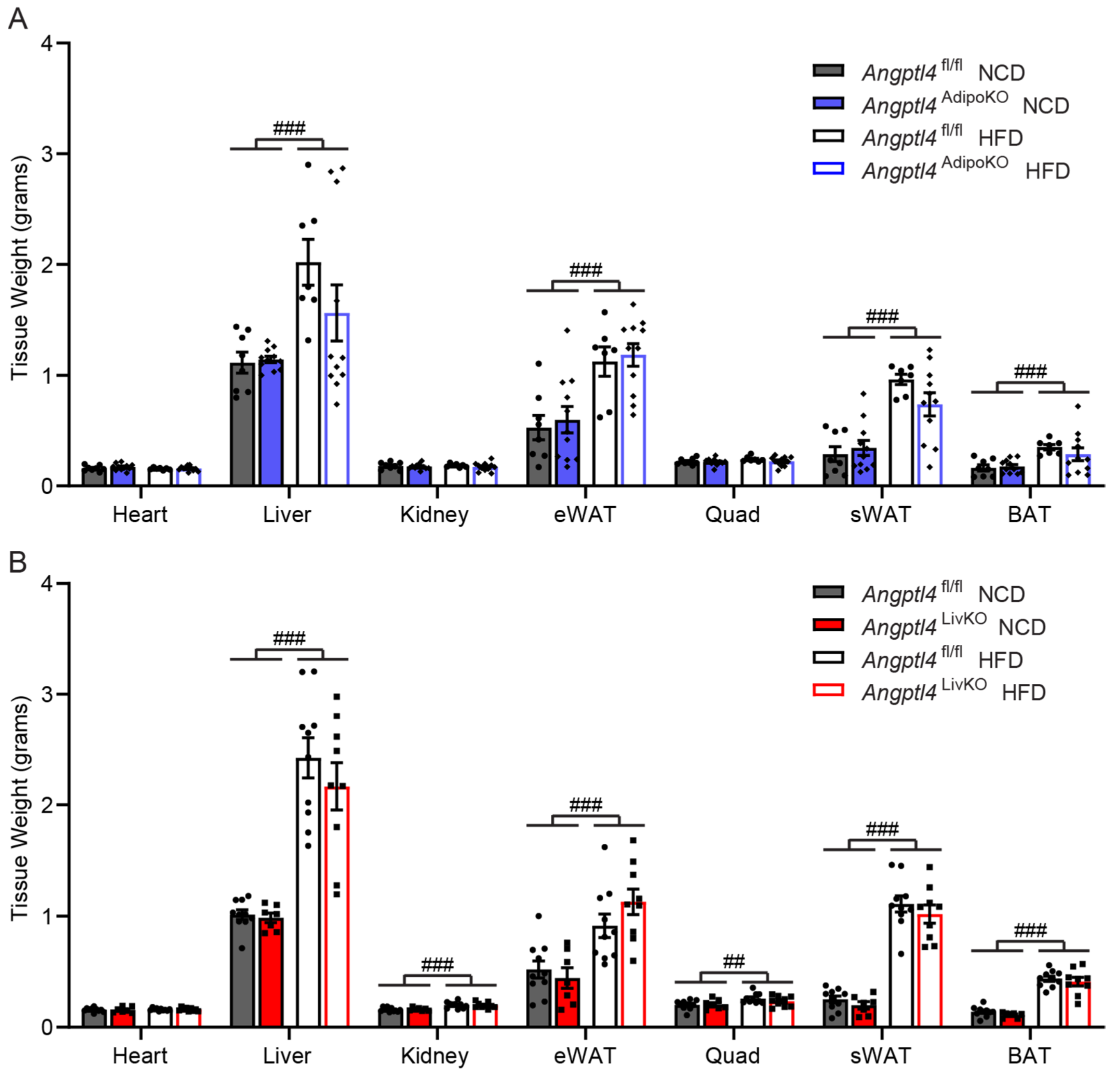


**Supplemental Figure 2: Tissue weights of *Angptl4*$^{AdipoKO}$ and *Angptl4*$^{LivKO}$ mice.** Tissues weights of heart, liver, kidney, epididymal white adipose (eWAT), quadriceps muscle (Quad), subcutaneous white adipose tissue (sWAT), and brown adipose tissue (BAT) in male *Angptl4*$^{AdipoKO}$ **(A)** and *Angptl4*$^{LivKO}$ **(B)** mice after 12 weeks on diet (mean±SEM; n=6-10/group). ###p<0.001 for dietary differences by two-way ANOVA.

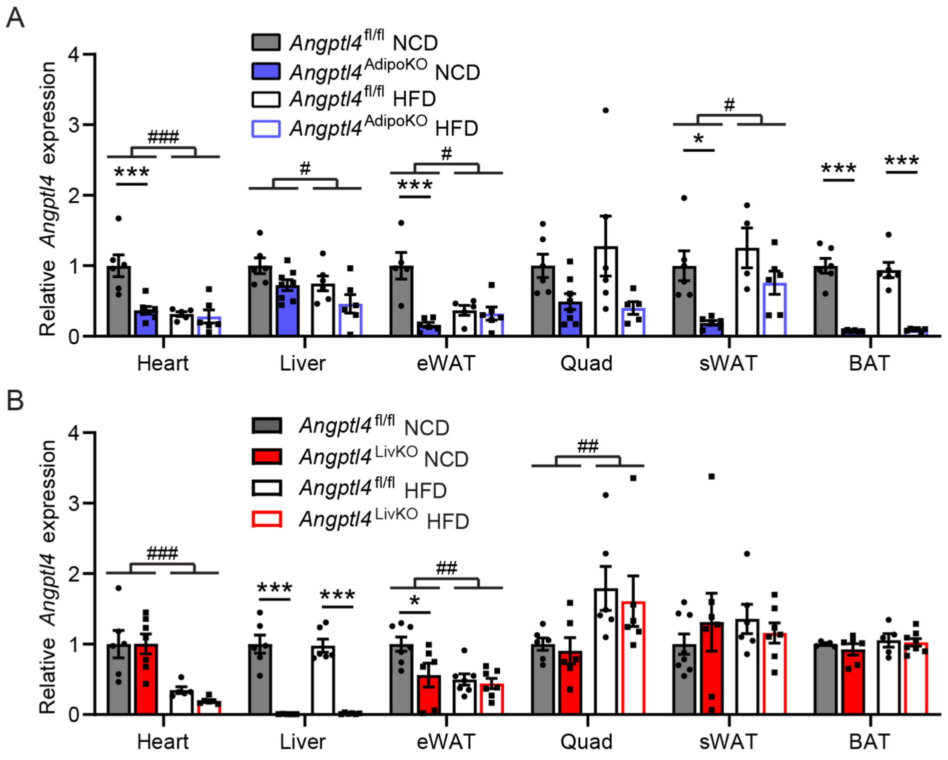


**Supplemental Figure 3: *Angptl4* expression in *Angptl4*$^{\mathrm{AdipoKO}}$ and *Angptl4*$^{\mathrm{LivKO}}$ mice fed a HFD for 12 weeks.** After 12 weeks on NCD or HFD, male *Angptl4*$^{\mathrm{fl/fl}}$, *Angptl4*$^{\mathrm{LivKO}}$, and *Angptl4*$^{\mathrm{AdipoKO}}$ mice were sacrificed and tissues were collected. mRNA expression of *Angptl4* from heart, liver, gonadal white adipose tissue (eWAT), quadriceps muscle (quad), subcutaneous white adipose tissue (sWAT), and brown adipose tissue (BAT) of fasted (6 h) male *Angptl4*$^{\mathrm{AdipoKO}}$ **(A)** and *Angptl4*$^{\mathrm{LivKO}}$ **(B)** mice (mean± SEM; n=4-8/group). #p<0.05, ##p<0.01, ###p<0.001 for dietary differences by two-way ANOVA. *p<0.05, **p<0.01, ***p<0.001 for individual genotype-specific differences by multiple comparison after two-way ANOVA (Tukey correction).

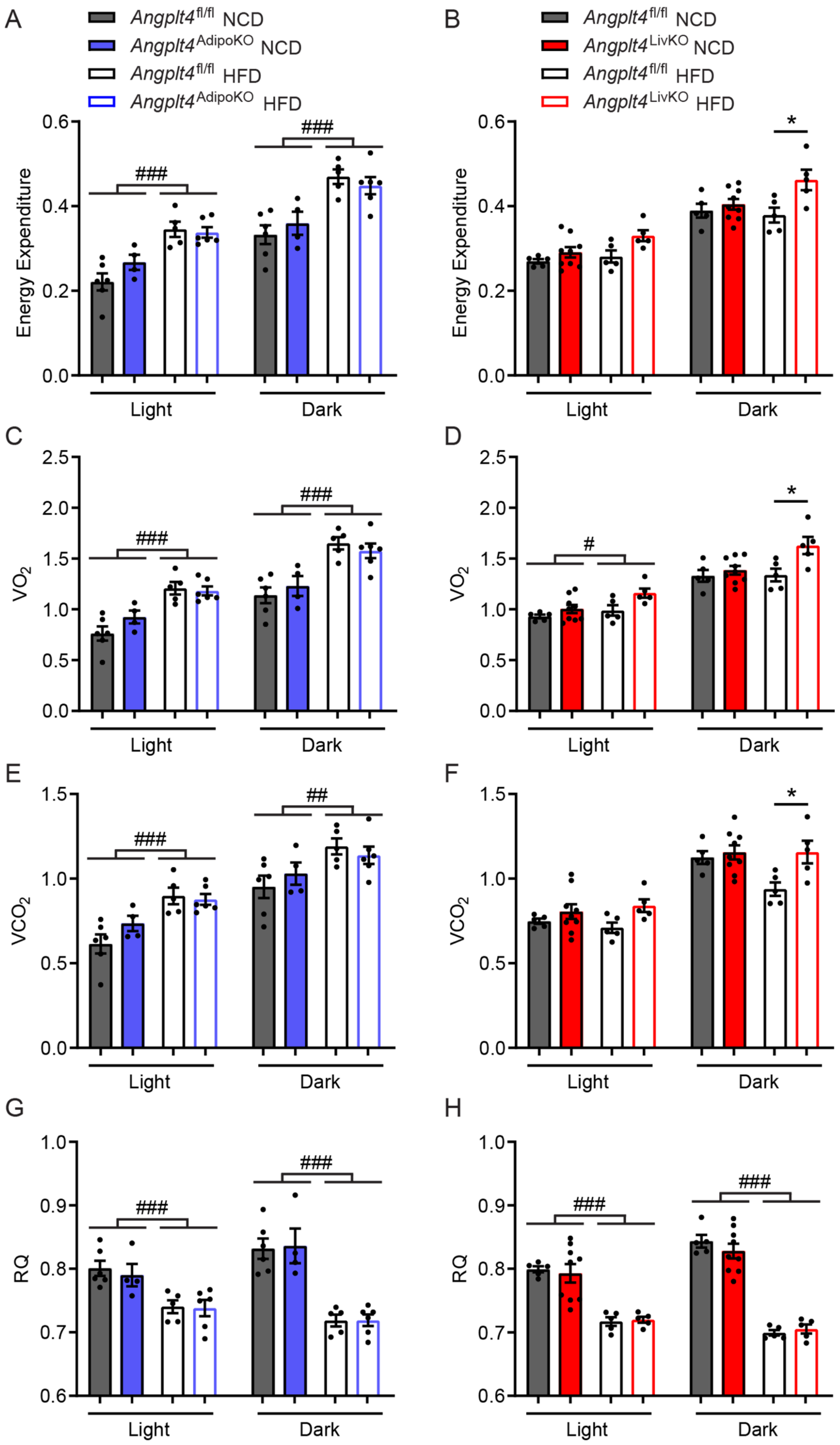


**Supplemental Figure 4. Respiratory measurements in male *Angptl4*[AdipoKO] and *Angptl4*[LivKO] mice.** After 12 weeks on NCD or HFD, male *Angptl4*[fl/fl], *Angptl4*[LivKO] and *Angptl4*[AdipoKO] mice were placed into metabolic cages. After 36 h of acclimation, energy expenditure **(A, B)**, $VO_2$ **(C, D)**, $VCO_2$ **(E, F)**, and respiratory quotient (RQ**, G, H)** were measured over 48 h (mean±SEM; n=4-9). Data are separated into the light (6 a.m- 6 p.m.) and dark (6 p.m.- 6 a.m.) cycles. #p<0.05, ##p<0.01, ###p<0.001 for dietary differences by two-way ANOVA. *p<0.05 for individual genotype-specific differences by multiple comparison after two-way ANOVA (Tukey correction).

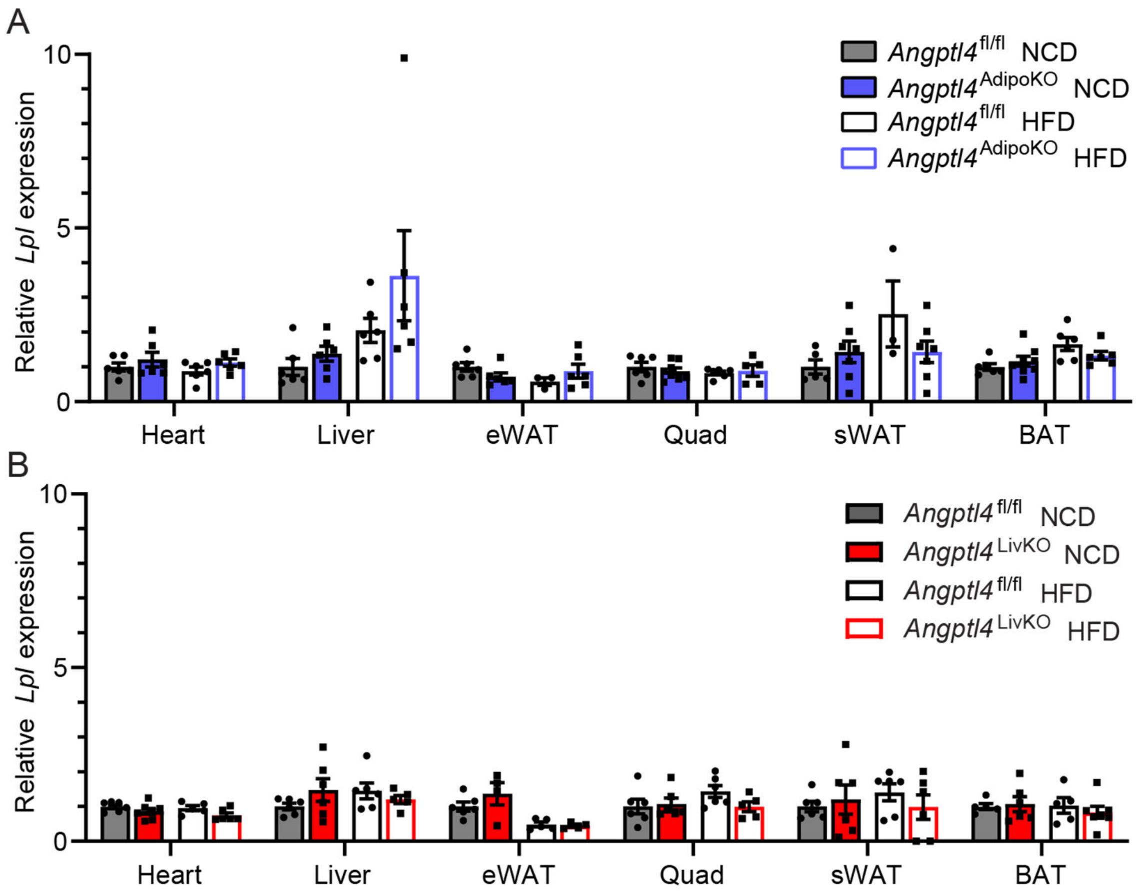


**Supplemental Figure 5: Lipoprotein Lipase expression in tissues from *Angptl4*$^{\mathbf{AdipoKO}}$ and *Angptl4*$^{\mathbf{LivKO}}$ mice.** Fasted (6 h) mRNA expression of *Lpl* in liver, heart, quadriceps muscle (quad), epididymal white adipose tissue (eWAT), subcutaneous white adipose tissue (sWAT), and brown adipose tissues (BAT) of male *Angptl4*$^{\text{AdipoKO}}$ **(A)** and *Angptl4*$^{\text{LivKO}}$ **(B)** mice fed either a normal chow diet (NCD) or a high fat diet (HFD; 60% by kCal) for 12 weeks starting at 8 weeks of age (mean±SEM; n=4-7/group).

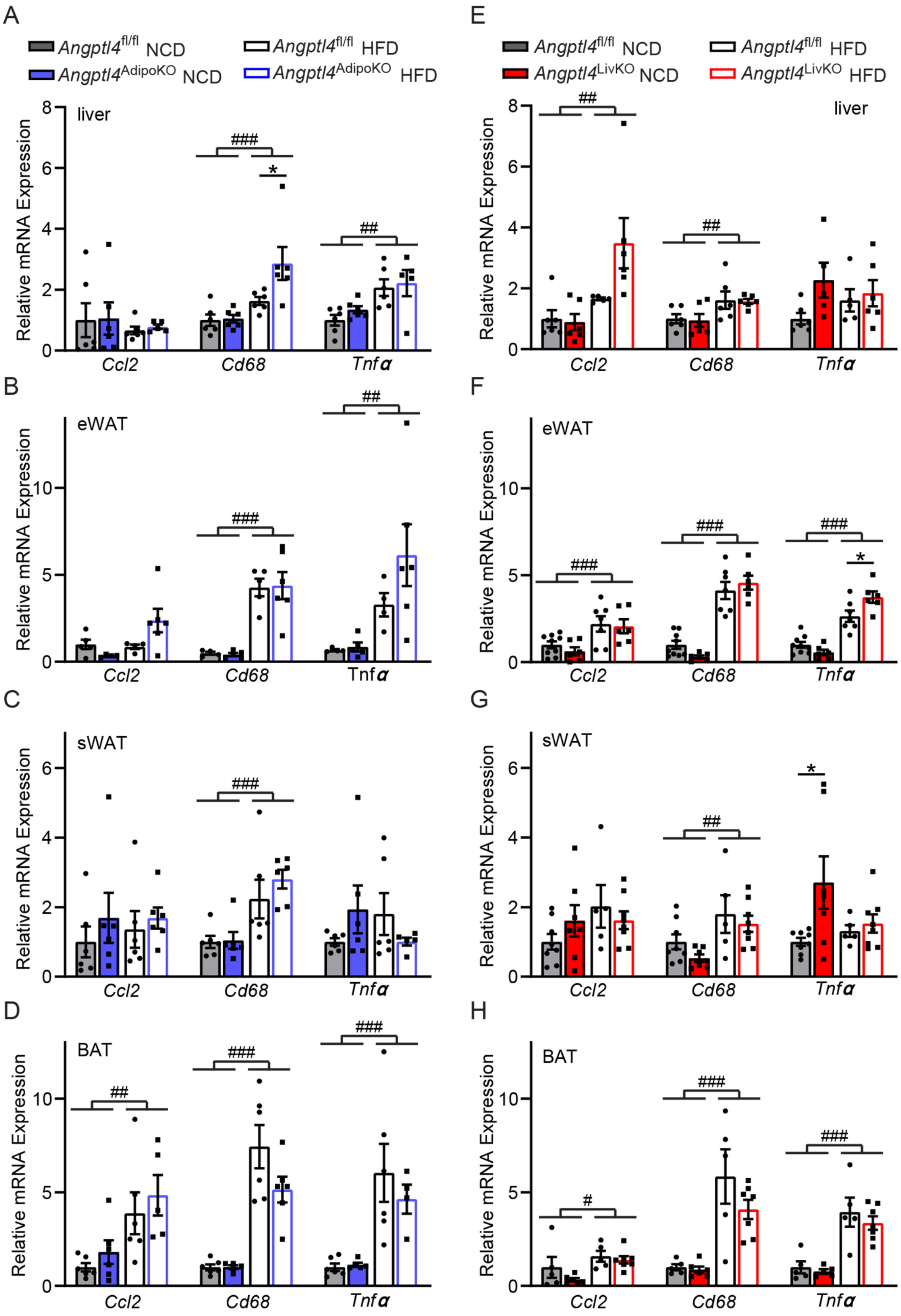


**Supplemental Figure 6: Inflammatory marker expression in tissues from *Angptl4*$^{AdipoKO}$ and *Angptl4*$^{LivKO}$ mice.** Fasted (6 h) mRNA expression of inflammatory markers *Ccl2*, *Cd68*, and *Tnf*α from liver tissue **(A and E)**, gonadal white adipose tissue (eWAT) **(B and F)**, subcutaneous white adipose tissue (sWAT)(**C and G**), and brown adipose tissues (BAT)**(D and H)** of male *Angptl4*$^{AdipoKO}$ **(A–D)** and *Angptl4*$^{LivKO}$ **(E–H)** mice fed either a normal chow diet (NCD) or a high fat diet (HFD; 60% by kCal) for 12 weeks starting at 8 weeks of age (mean±SEM, n=4-7/group). #p<0.05, ##p<0.01, ###p<0.001 for dietary differences by two-way ANOVA. *p<0.05, for individual genotype-specific differences by multiple comparison after two-way ANOVA (Tukey correction).

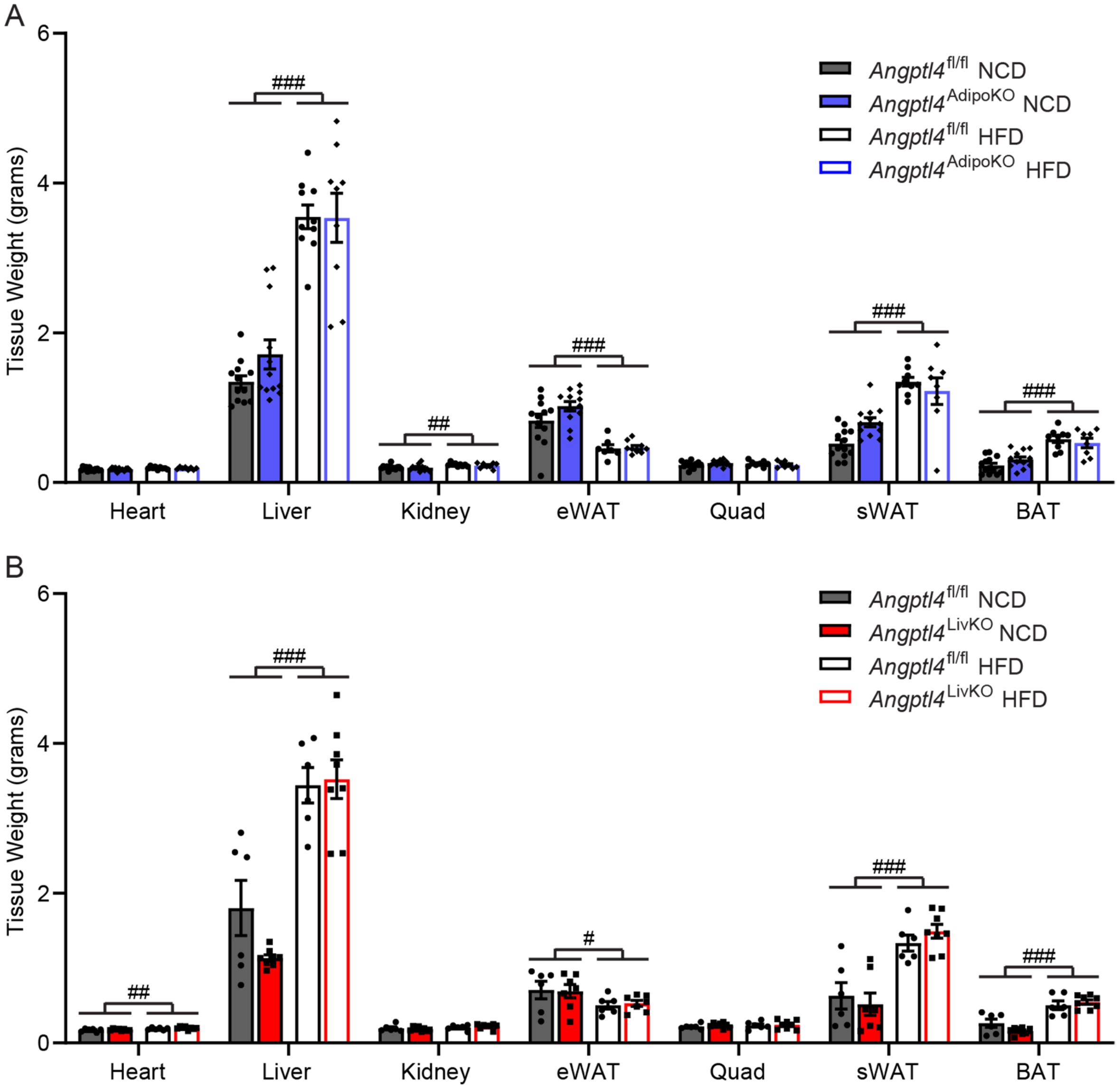


**Supplemental Figure 7: Tissues weights from *Angptl4*$^{\text{AdipoKO}}$ and *Angptl4*$^{\text{LivKO}}$ mice after chronic high-fat feeding.** Tissues weights of heart, liver, kidney, epididymal white adipose (eWAT), quadriceps muscle (Quad), subcutaneous white adipose tissue (sWAT), and brown adipose tissue (BAT) in male *Angptl4*$^{\text{AdipoKO}}$ **(A)** and *Angptl4*$^{\text{LivKO}}$ **(B)** mice after 6 months of either normal chow diet (NCD) or high fat diet (HFD) feeding (mean± SEM; n=6-12/group). #p<0.05, ##p<0.01, ###p<0.001 for dietary differences by two-way ANOVA.

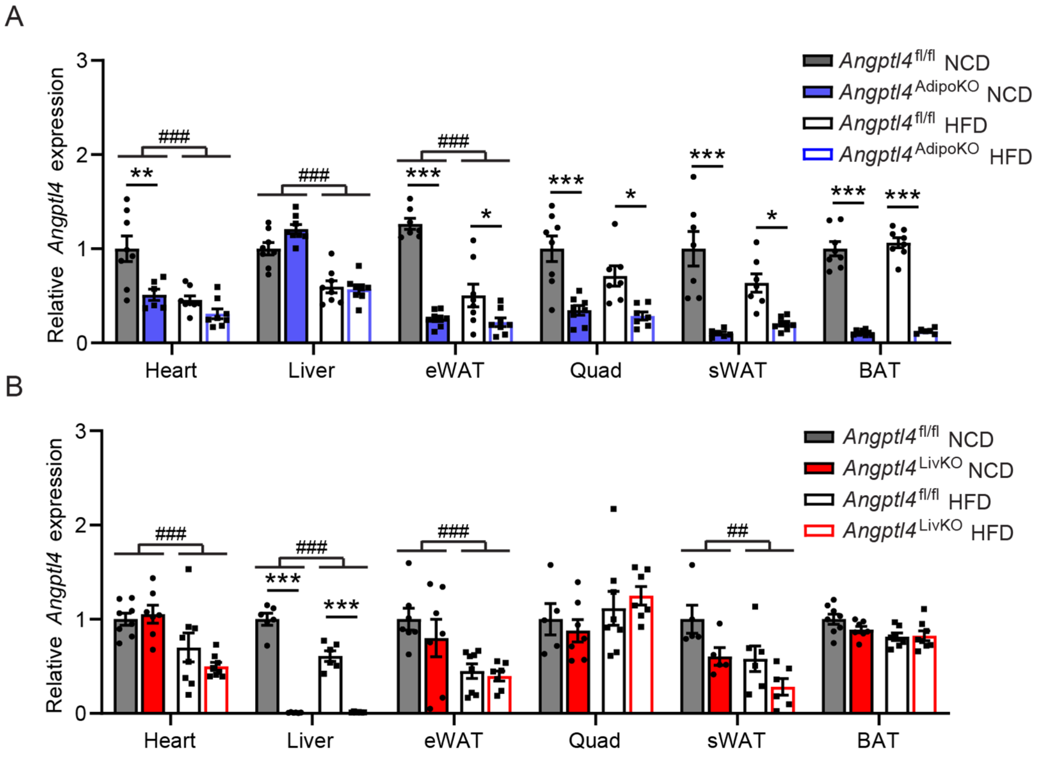


**Supplemental Figure 8: *Angptl4* expression in *Angptl4*$^{AdipoKO}$ and *Angptl4*$^{LivKO}$ mice after chronic high-fat feeding.** mRNA expression of *Angptl4* from heart, liver, gonadal white adipose tissues (eWAT), quadriceps muscle (quad), subcutaneous white adipose tissue (sWAT), and brown adipose tissues (BAT) of fasted (6 h) male *Angptl4*$^{AdipoKO}$ **(A)** and *Angptl4*$^{LivKO}$ ***(*B)** mice after 6 months on NCD or HFD (mean± SEM; n=5-8/group). ##p<0.01, ###p<0.001 for dietary differences by two-way ANOVA. *p<0.05,**p<0.01,***p<0.001 for individual genotype-specific differences by multiple comparison after two-way ANOVA (Tukey correction).

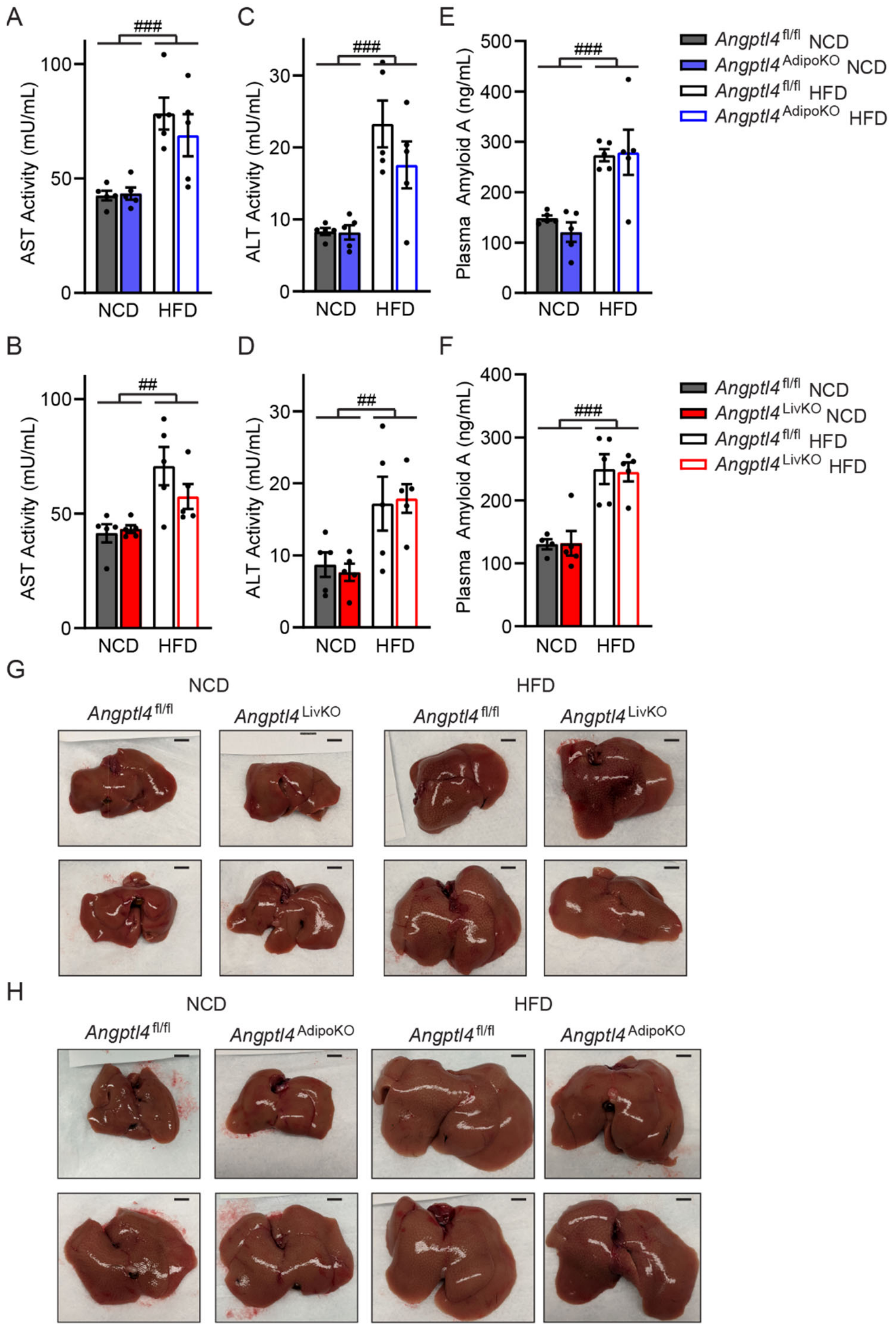


**Supplemental Figure 9: Liver phenotypic markers in *Angptl4*$^{AdipoKO}$ and *Angptl4*$^{LivKO}$ mice after chronic high-fat feeding.** AST activity (**A,B**), ALT activity (**C,D**) and Amyloid A levels (**E,F**) from the plasma of fasted (6 h) male *Angptl4*$^{AdipoKO}$ **(A,C,E)** and *Angptl4*$^{LivKO}$ **(B,D,F)** mice fed either a NCD or HFD for 6 months (mean±SEM, n=5-6/group). ##$p<0.01$, ###$p<0.001$ for dietary differences by two-way ANOVA. Representative pictures of livers from male *Angptl4*$^{LivKO}$ **(G)** or *Angptl4*$^{AdipoKO}$ **(H)** mice after 6 months of either NCD or HFD feeding.

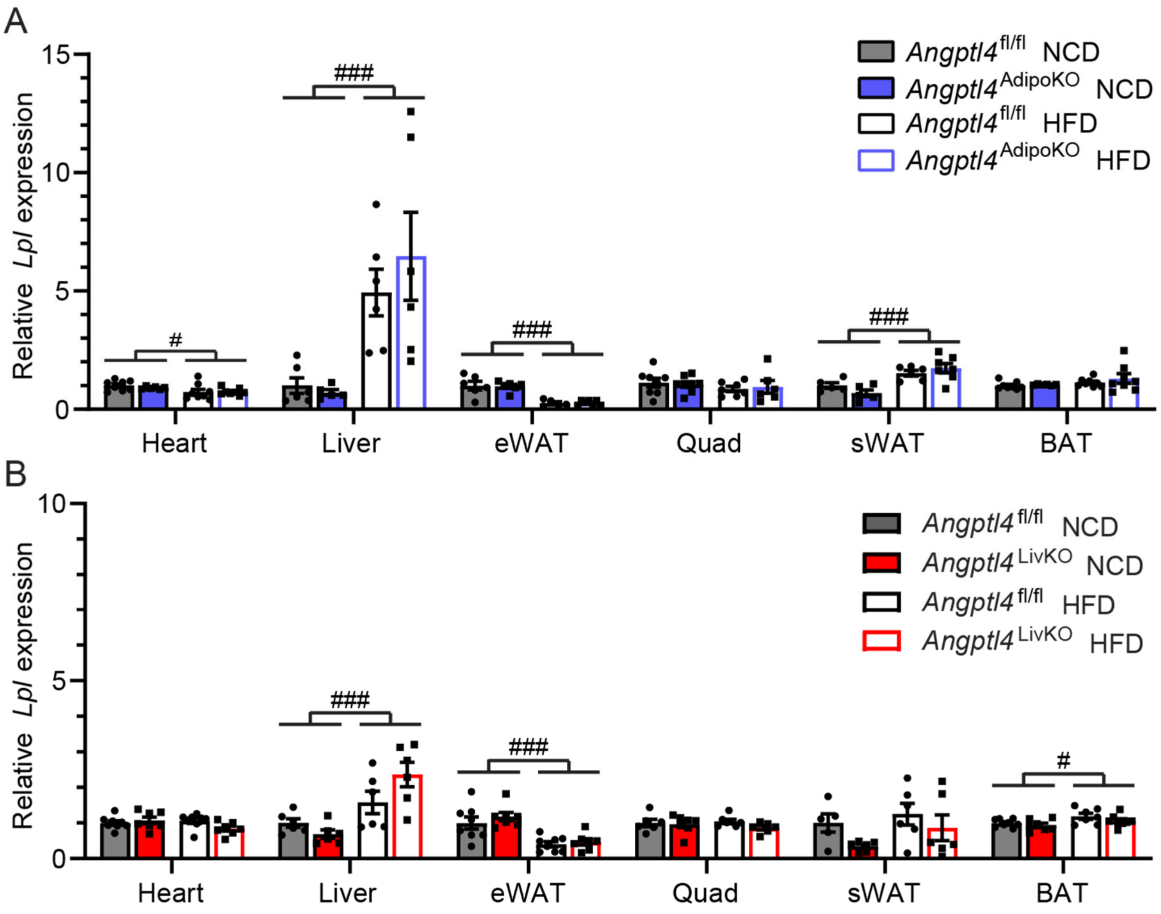


**Supplemental Figure 10: Lipoprotein lipase expression in tissues from *Angptl4*$^{\mathbf{AdipoKO}}$ and *Angptl4*$^{\mathbf{LivKO}}$ mice after chronic high-fat feeding.** Fasted (6 h) mRNA expression of *Lpl* in liver, heart, quadriceps muscle (quad), epididymal white adipose tissue (eWAT), subcutaneous white adipose tissue (sWAT), and brown adipose tissues (BAT) of male *Angptl4*$^{AdipoKO}$ **(A)** and *Angptl4*$^{LivKO}$ **(B)** mice after 6 months of either normal chow diet (NCD) or a high fat diet (HFD) feeding (mean±SEM; n=4-7/group). #p<0.05, ##p<0.01, ###p<0.001 for dietary differences by two-way.

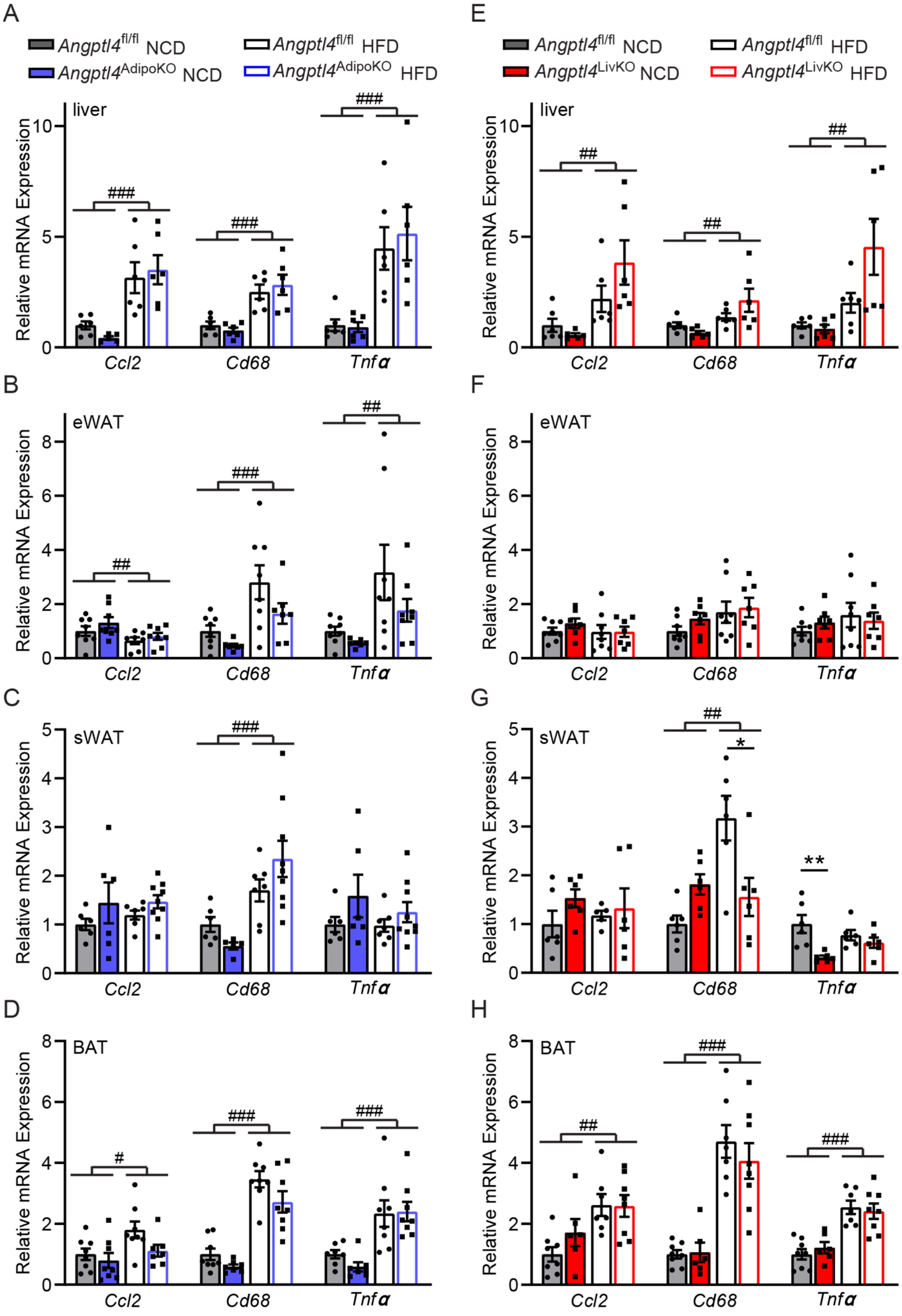


**Supplemental Figure 11: Inflammatory marker expression from *Angptl4*$^{AdipoKO}$ and *Angptl4*$^{LivKO}$ mice after chronic high-fat feeding.** Fasted (6 h) mRNA expression of inflammatory markers *Ccl2*, *Cd68*, and *Tnf*α from liver tissue **(A and E)**, gonadal white adipose tissue (eWAT)**(B and F)**, subcutaneous white adipose tissue (sWAT)**(C and G)**, and brown adipose tissues (BAT)**(D and H)** of male *Angptl4*$^{AdipoKO}$ **(A–D)** and *Angptl4*$^{LivKO}$ **(E–H)** mice fed either a normal chow diet (NCD) or a high fat diet (HFD; 60% by kCal) for 6 months (mean±SEM, n=7-8/group). #p<0.05, ##p<0.01, ###p<0.001 for dietary differences by two-way ANOVA. *p<0.05, **p<0.01 for individual genotype-specific differences by multiple comparison after two-way ANOVA (Tukey correction).

**Supplemental Table 1: Primers A-F used in Supplemental Figure 1**

| Primer | Forward | Reverse |
|---|---|---|
| **A** | GCTGCCCTGGTGCTATG | TGAGCCAGCAAGTTCATCTC |
| **B** | TTTGCAGACTCAGCTCAAGG | TCCATTGTCTAGGTGCGTGG |
| **C** | TTTGCAGACTCAGCTCAAGG | TGTGTAAGTGGGTGGCGTTGGG |
| **D** | GACTCAGCTCAAGGCTCAAA | TCTGGCTCTGAAGATTCTGTATTC |
| **E** | ATGACTTCAGATGGAGGCTGG | AATTGGCTTCCTCGGTTCCC |
| **F** | CAACTAGCTGGGCCCTTAAT | ATCCACAGCACCTACAACAG |
